# The Speech Reception Threshold Can be Estimated Using EEG Electrodes In and Around the Ear

**DOI:** 10.1101/2024.12.02.625819

**Authors:** Heidi B. Borges, Johannes Zaar, Emina Alickovic, Christian B. Christensen, Preben Kidmose

## Abstract

**Abstract:** *Objective:* Previous studies have demonstrated that the speech reception threshold (SRT) can be estimated using scalp electroencephalography (EEG), referred to as SRT_neuro._ The present study assesses the feasibility of using ear-EEG, which allows for discreet measurement of neural activity from in and around the ear, to estimate the SRT_neuro_. Such an estimate can be highly useful e.g. for continuously adjusting noise-reduction algorithms in hearing aids or for logging the SRT in the user’s natural environment.

*Approach:* Twenty young normal-hearing participants listened to audiobook excerpts at varying signal-to-noise ratios (SNRs) whilst wearing a 66-channel EEG cap and 12 ear-EEG electrodes. A linear decoder was trained on different electrode configurations to estimate the envelope of the audio excerpts from the EEG recordings. The reconstruction accuracy was determined by calculating the Pearson’s correlation between the actual and the estimated envelope. A sigmoid function was then fitted to the reconstruction-accuracy-vs-SNR data points, with the midpoint of the sigmoid serving as the SRT_neuro_ estimate for each participant.

*Main results:* Using only in-ear electrodes, the estimated SRT_neuro_ was within 3 dB of the behaviorally measured SRT (SRT_beh_) for 6 out of 20 participants (30%). With electrodes placed both in and around the ear, the SRT_neuro_ was within 3 dB of the SRT_beh_ for 19 out of 20 participants (95%) and thus on par with the reference estimate obtained from full-scalp EEG. Using only electrodes in and around the ear from the right side of the head, the SRT_neuro_ remained within 3 dB of the SRT_beh_ for 19 out of 20 participants.

*Significance:* These findings suggest that the SRT_neuro_ can be reliably estimated using ear-EEG, especially when combining in-ear electrodes and around-the-ear electrodes.

## 1 Introduction

Many people with hearing impairment struggle to understand speech in noisy environments, even when using hearing aids that compensate for their hearing loss [1]. They describe difficulty in understanding speech in noise as the most limiting aspect of being hearing impaired [2]. The ability to understand speech in noisy conditions is often assessed using the speech reception threshold (SRT). Measuring SRT within a hearing aid could enable optimized noise reduction, applying noise reduction targeting increased speech intelligibility when intelligibility is low, and applying less noise reduction to preserve the naturalness of sound when intelligibility is high. An ongoing measure of the SRT could also be highly useful to monitor a user’s speech intelligibility challenges in their natural environment, or as an at home SRT assessment. Traditionally, the SRT procedure determines the signal-to-noise ratio (SNR) at which a person can repeat 50% of the speech presented to them. This method has limitations due to the reliance on the participant’s responses and is therefore not suitable for the purposes described above.

Recent studies have successfully estimated the SRT from scalp electroencephalography (EEG) in normal-hearing listeners [3–5]. The SRT was estimated from scalp EEG by calculating the reconstruction accuracy of a linear decoder reconstructing the speech envelope from EEG and afterwards fitting a sigmoid function to the reconstruction-accuracy-vs-SNR data points [3–5]. Although this approach eliminates the need for the participant to respond, scalp EEG is not feasible for regular daily use and the method is therefore not suitable for integration into a hearing aid or other hearables.

Compared to traditional scalp EEG, ear-EEG [6,7] offers a more discreet and mobile measurement setup with EEG electrodes placed in or around the ear, enabling EEG recording in everyday life. While speech intelligibility has been estimated from scalp EEG before [3–5] in young normal hearing listeners, to the authors knowledge, estimation of the SRT from electrodes in and around the ear has not been conducted previously. Other related measures have been investigated using ear-EEG, such as estimation of pure-tone thresholds using classic Auditory Steady-State Response stimuli in populations of normal hearing and hearing impairment [8,9] and natural sounds [10], as well as auditory attention decoding using linear decoding methods on a population of normal hearing [11,12]. Mirkovic et al (2016) [12] used ear-EEG recordings to classify the attended speaker in a 2-talker scenario in a young normal hearing population. They demonstrated that changes in the accuracy of the reconstructed speech envelope related to the auditory attention could be measured from electrodes placed around the ear. Moreover, Fiedler et al. (2017) [12] recorded EEG from the left ear canal with a FT7 reference and used an encoding model to predict the EEG signals from the speech envelope to classify the attended speaker in a two-talker selective attention scenario. Collectively, the results of these studies demonstrate that the encoding of the speech envelope in the auditory system can be measured using ear-EEG.

In the present study, we investigate whether the SRT can be estimated from EEG recorded from electrodes placed in and around the ear in a young, normal-hearing population. This work extends the study by Borges et al. (2024) [5], where a linear decoder was trained to predict the envelope of the audio stimuli. Reconstruction accuracy was first assessed by calculating the Pearson’s correlation between the actual and predicted envelopes. A sigmoid function was then fitted to the reconstruction accuracy versus SNR data, with the midpoint of the sigmoid serving as the SRT_neuro_ estimate. The SRT_neuro_ was found to be within 3 dB of the behaviorally measured SRT (SRT_beh_) for all participants. The present study is based on the same methodology and data as described in Borges et al (2024) [5] but focusing specifically on estimating the SRT_neuro_ from the ear-EEG data, which was not included in the original study. This approach enables a direct comparison between the performance of the two studies.

## 2 Materials and methods

The data analyzed was acquired in a previous study investigating estimation of a behaviorally determined SRT (SRT_beh_) from scalp EEG using a linear decoder to reconstruct the envelope of the presented audiobook excerpts [5].

### 2.1 Participants

Participants were included based on self-reported normal hearing, age between 18 and 30 years, and Danish as a native language. The normal hearing criterion was confirmed through audiometric testing across frequencies of 125 Hz – 8 kHz, allowing a maximum hearing loss of 20 dB, which is considered normal hearing according to WHO guidelines [13]. The study was approved by the Institutional Review Board at Aarhus University with the approval number TECH-2022-004.

### 2.2 Experimental setup

#### EEG

All EEG data were recorded at a sampling rate of 4096 Hz with an average reference. Scalp EEG was recorded using the Biosemi Active EEG (Amsterdam, Netherlands) system with a 64-electrode cap and 2 additional mastoid electrodes. The 12 in-ear EEG electrodes and an additional electrode at Fpz were recorded using the SAGA32+/64+ system (TMSi, Oldenzaal, Netherlands). The 12 in-ear EEG electrodes had a diameter of 4 mm and were made of silver/silver-chloride (Ag/AgCl) [14]. The in-ear EEG electrodes were placed on individually modelled earpieces in positions ExA, ExB, ExC, ExT, ExI and ExK where “x” will be replaced with “L” if placed in the left ear and with “R” if placed in the right ear, see figure 1(a), for the location of the scalp electrodes see figure 1(b). Placement and manufacturing of the earpiece was conducted as described in Kidmose et al (2013) [15]. The ear-EEG earpiece was modelled from the individual ear-impressions using a CAD software (EarMouldDesigner, 3Shape, Denmark), and was made from silicone with shore 60 (Detax softwear 2.0, Detax GmbH, Germany). Before inserting the earpiece, the participants’ ears were cleaned with a cotton swap soaked in water. The quality of the ear-EEG signal was assessed in a live viewer and the resemblance to EEG and the presence of the typical signatures of eyeblinks and jaw clenches in the EEG signal was assessed. If the signatures were not present, adding more water to the ear and/or taking the ear-EEG earpiece in and out was performed.

**Figure 1.**
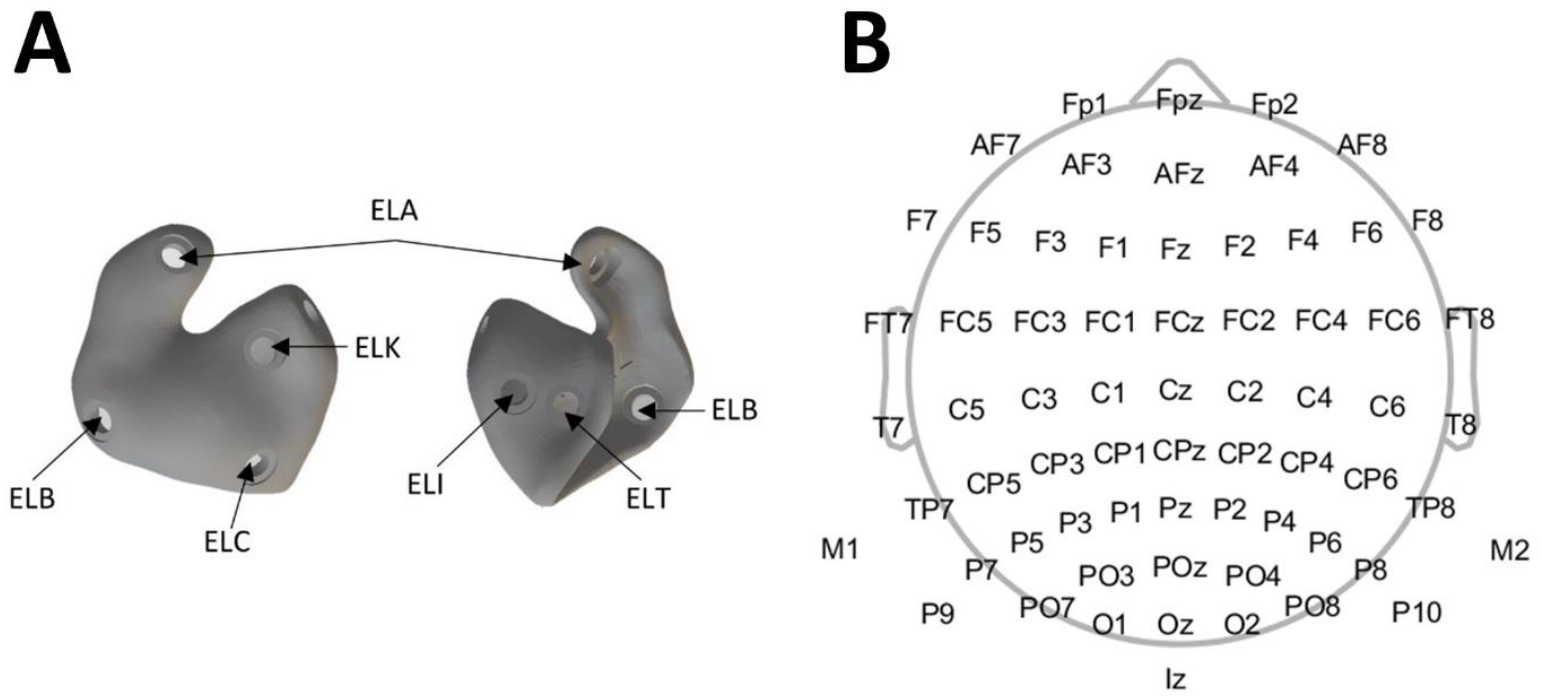
Panel A: an example of a left-side ear-EEG earpiece and the location of the used electrodes. Panel B: the location of the scalp electrodes.

#### Stimulus presentation

During the SRT_beh_ estimation, the stimuli were presented using Etymotic ER-1 insert Earphones (Etymotic Research, Inc., IL, USA) through disposable foam ear tips, via a soundcard (RME hammerfall DSB multiface II, Audio AG, Germany). For the EEG experiment, the stimuli were presented in a similar manner, except that the sound was delivered through the sound bore of the ear-EEG earpiece instead of the foam tips.

#### Behavioral SRT

The SRT_beh_ was measured at 50% words correct using the Danish Hearing-In-Noise Test (HINT) sentences presented in stationary speech-shaped noise, using an adaptive threshold tracking procedure, as described in Borges et al. 2024 [5].

#### An Estimated SRT from EEG

To estimate the SRT from EEG (SRT_neuro_), audiobook excerpts of approximately 60 seconds in duration, trimmed in duration to preserve complete sentences, were played to participants across six conditions, including five different signal-to-noise ratios (SNRs) and a clean speech condition. The speech level was set at 65 dB SPL and different levels of stationary speech-shaped noise were added for the SNR conditions. The SNR conditions were defined relative to the subject-specific SRT_beh_ as: SRT_beh_ -4dB, SRT_beh_ -2dB, SRT_beh_, SRT_beh_ +2dB, SRT_beh_ +4dB. The audiobook material was lowpass-filtered using a first-order lowpass filter with a cutoff frequency of 2 kHz. This was done to approximate the third-octave power spectral density of the HINT speech corpus. A total of 16 audiobook excerpts were presented for each condition (96 in total) in a randomized design, as described in Borges et al. 2024 [5].

### 2.3 Analysis

#### Stimuli

The envelope of the stimuli was extracted by computing the absolute value of the analytic signal, derived using the Hilbert transform, for each audiobook excerpt. To compensate for nonlinearities in the auditory system, power-law compression with an exponent of 0.6 was applied, as described in Biesmans et al. (2017) [16]. The signal was then bandpass-filtered between 1 and 8 Hz and down-sampled to 64 Hz, following the procedure outlined in Borges et al. 2024 [5].

#### Preprocessing and cleaning of EEG data

Initially, the ear and scalp electrodes were processed separately. Ear-EEG electrodes were average-referenced and scalp electrodes Fpz-referenced. Electrodes yielding signals that were flat or had a standard deviation (SD) greater than three times the mean SD of all other electrode signals were rejected. The SD was calculated after highpass-filtering with a 1-Hz cutoff third-order highpass filter. The rejected scalp electrodes were replaced using spherical interpolation. The rejected ear electrodes were either omitted (AEar configurations) or replaced by the average of local non-rejected ear electrodes (Ear configurations, see details in the electrode configurations paragraph). The scalp and ear electrodes were then referenced to the common measurement electrode (Fpz), and the electrodes sets were merged. The channels were down-sampled to 256 Hz to lower computation time. The channels were then bandpass-filtered between 1 and 8 Hz using a third-order bandpass filter by means of the “filtfilt” function in Matlab (The MathWorks Inc), ultimately a 12^th^ order filter is applied due to the application method. Afterwards the channels were resampled to 64 Hz and epoched. To handle small differences in sampling frequency between the amplifiers, the length of the trials in the ear-EEG data, marked by a start and end trigger, was used to determine the trial length in the scalp data. To conduct artefact rejection, extreme values were removed. This was done by referencing the ear-EEG electrodes to an average ear-EEG reference and the scalp electrodes to Cz. Then all values greater than 100 μV and smaller than −100 μVwere removed along with 32 datapoints before and after. The gap was then filled using autoregressive modelling, which was implemented using the “fillgaps” function in Matlab.

#### Electrode configurations

To assess the feasibility of using ear-EEG to estimate the SRT_neuro_, 13 different reference configurations using ear-EEG electrodes and electrodes around the ear (T7/8 and M1/2) were investigated (Figure 2). In addition, a scalp-EEG dataset (Scalp), where 64 scalp electrodes were referenced to the average of the mastoids, was included to account for small differences in the EEG preprocessing when comparing the present results to those reported in Borges et al. 2024 [5]. After referencing, all channels were normalized by dividing each channel by its own SD. The normalization was performed with the SD of each individual channel to attenuate potentially noisy channels.

**Figure 2.**
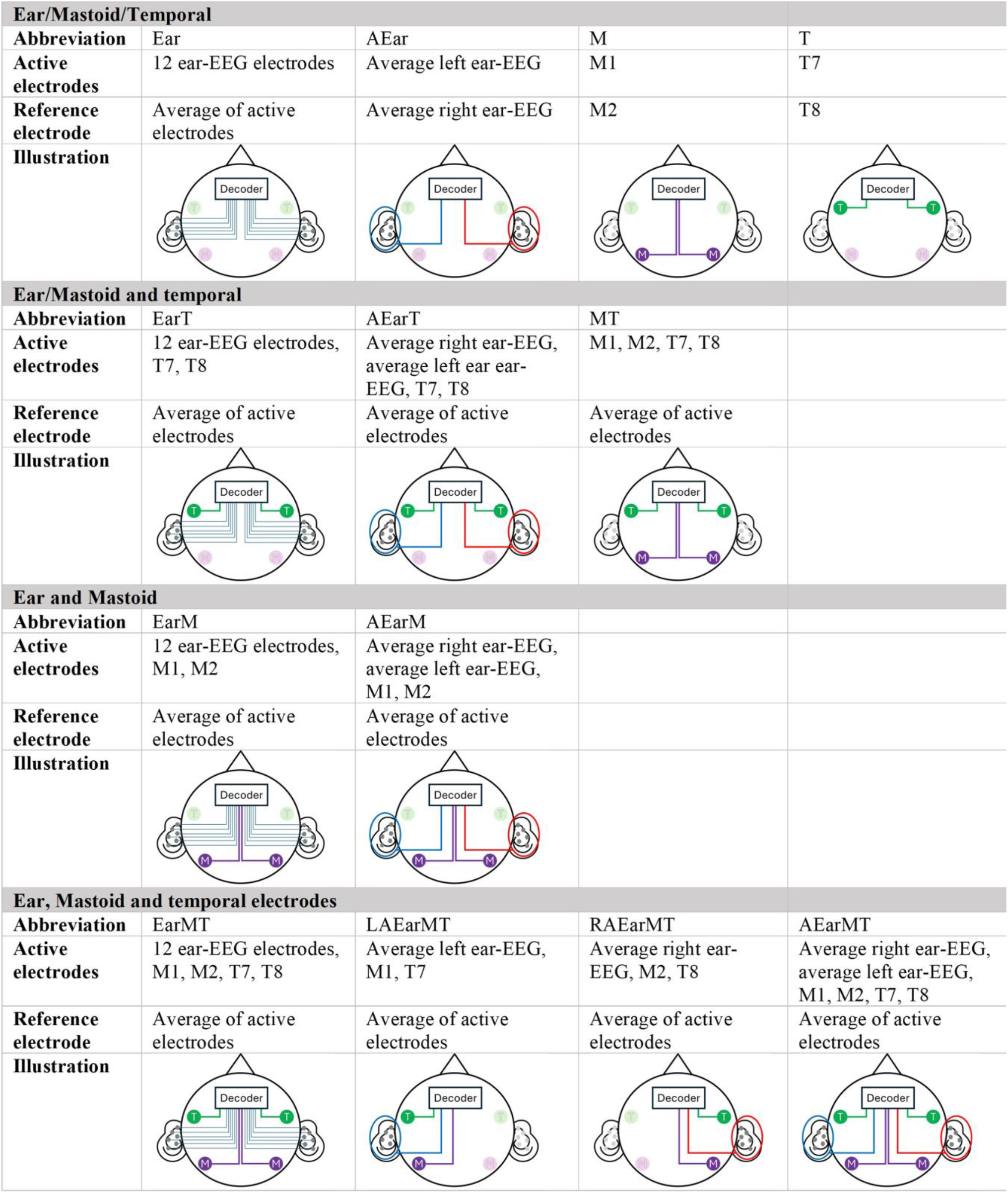
The different electrode configurations using electrodes in and around the ear as input for the decoder. Each line going into the decoder is a channel input. The abbreviation for the electrode configuration is shown under “Configuration”, with the active electrodes and their corresponding references described.

#### Decoder

A linear decoder was trained to reconstruct the speech envelope of the audio excerpts from the EEG signals, following the approach described in Borges et al. 2024 [5]. This was done separately for each electrode configuration, using only the clean speech condition for training, and testing on the conditions with noise (SRT_beh_-4dB, SRT_beh_-2dB, SRT_beh_, SRT_beh_+2dB, SRT_beh_+4dB). The decoder was estimated using regularized linear regression [17,18], with a decision window of [-100, 350] and a regularization parameter (λ) range of [10^−4^, 10^10^]. Reconstruction accuracy was calculated as Pearson’s correlation between the reconstructed envelope and the original speech envelope. To obtain the reconstruction accuracy for clean speech, the decoder was trained using a leave-one-trial-out approach: training on all but one of the clean speech excerpts, calculating the decoder accuracy for the left-out excerpt, repeating the procedure for all clean speech excerpts and then averaging the reconstruction accuracies across all excerpts. Reconstruction accuracy for the noise floor was obtained by calculating the Pearson’s correlation between the reconstructed envelope from each audio excerpt with noise (80 excerpts) and 68 mismatched envelopes from the same audiobook, which underwent the same preprocessing but were not used as experimental stimuli, as described in Borges et al. 2024 [5].

#### Estimating the SRT value

To estimate the SRT_neuro_ for each participant, a sigmoid function was fitted to the reconstruction-accuracy-vs-SNR data, as described in Borges et al. 2024 [5]. The sigmoid function as described by Farris-Trimble and McMurray (2013) [19] was used, see equation 1.

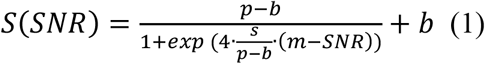

In equation 1 p represents the mean reconstruction accuracy of the clean speech condition, b is the reconstruction-accuracy noise floor, s denotes the slope and m the midpoint-value. The midpoint-value was used as the SRT_beh_ estimate. For this approach to be sound, an increase of reconstruction accuracy with increasing SNR is needed. Therefore, a permutation test was performed, assessing whether the reconstruction accuracies obtained for the condition’s clean speech, SRT_beh_+2dB and SRT_beh_+4dB were higher than those obtained for the conditions SRT_beh_-2dB, SRT_beh_-4dB and the noise floor. The permutation test had a precision of 0.01 and a significance level of 5% (α=0.05). The above described sigmoid fit was only performed if (i) the permutation test showed a significant increase, (ii) the average reconstruction accuracy for a condition was above the average clean speech reconstruction accuracy for no more than two SNR conditions, and (iii) the average reconstruction accuracy was below the noise floor average reconstruction accuracy for no more than two conditions. The sigmoid fit was performed 100 times with a random initialization point for the m value ±10 dB from -2.52 dB, which was the SRT_beh_ for normal-hearing participants in the HINT validation study [20]. The s parameter was restricted to be non-negative, and if the SRT_neuro_ was outside the range of [-40, 40] dB, the fit was rejected due to an unrealistic estimation. The 10 best fits were identified based on highest r-squared value and the final parameters were obtained as the mean of the respective parameters across these 10 best fits. An example of a fit from a representative subject with the ear electrode configuration (Ear) is shown in figure 3.

**Figure 3.**
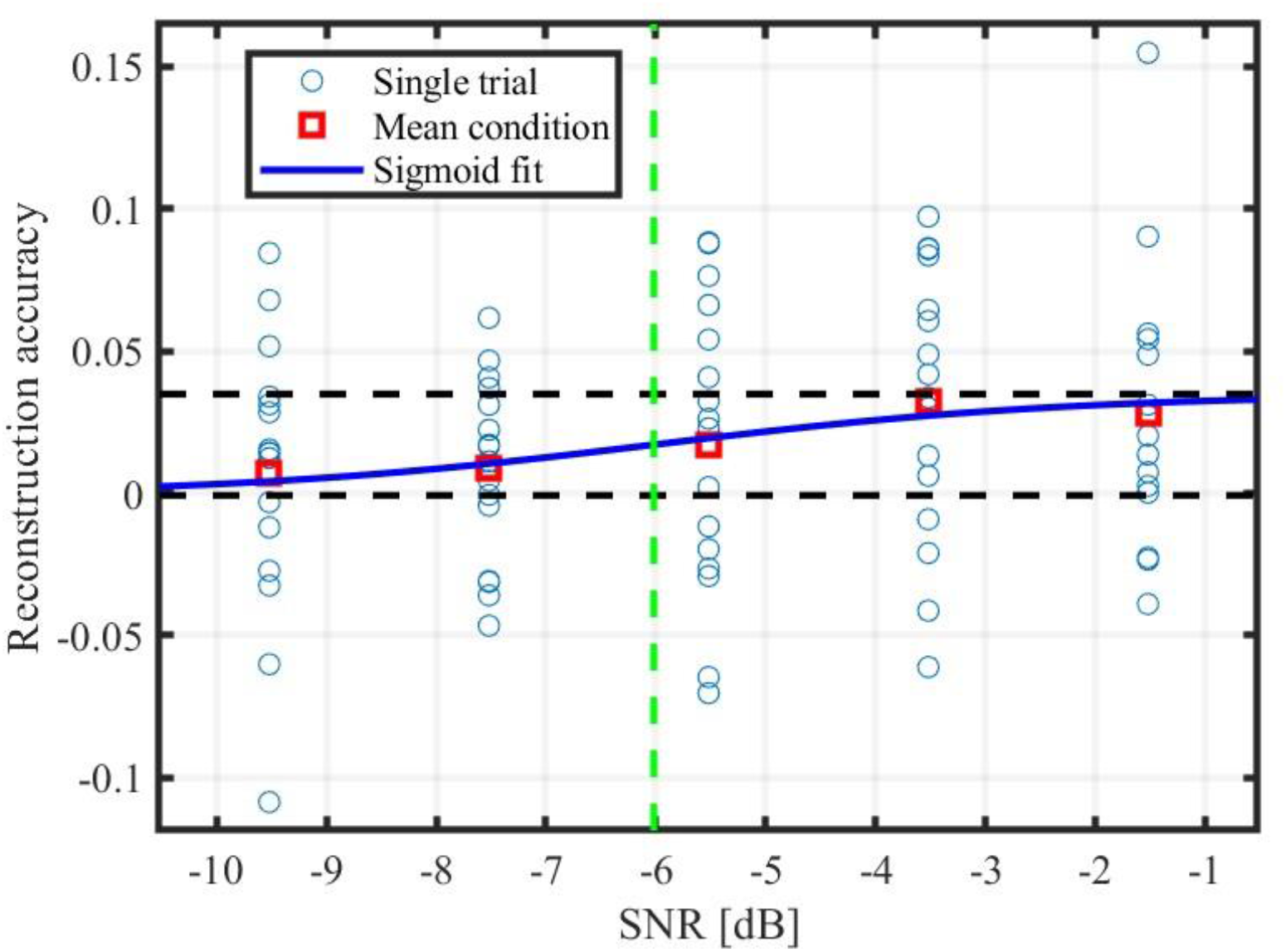
Example of a sigmoid fit to single-subject data from an ear electrode configuration. Each circle shows the reconstruction accuracy for one single trial. The across-trial means for each condition are shown as red squares. The maximum and minimum values used in the sigmoid fit (p and b parameters) are shown with dotted black lines the estimated SRT_neuro_ with a vertical dotted green line and the sigmoid fit is shown in dark blue.

#### Linear mixed-effects model

To investigate the relationship between SNR and reconstruction accuracy for each of the electrode configurations, a linear mixed-effects model was fitted in R using the nlme library [21,22] and maximum likelihood criteria. Participant number (P) was modeled as a random effect, allowing for an individual offset, SNR was modeled as a fixed effect and reconstruction accuracy (R) as the dependent variable resulting in the model: R ∼ SNR + (P|1). Residuals were analyzed for normality by observing the qq-plot and histogram of residuals. Statistical significance of the fixed effect was assessed with a univariate Wald test with α=0.05.

#### Comparison of the variance of difference between the SRT_beh_ and the SRT_neuro_

A chi-square test was performed to test whether the variance of the difference between the SRT_beh_ and the SRT_neuro_ for the configurations with electrodes in and around the ear differed from the scalp electrode configuration, which served as a reference. The null hypothesis was that the variance for each of the electrode configurations with electrodes in and around the ear comes from a normal distribution with the same variance as the difference between the SRT_beh_ and the SRT_neuro_ in the Scalp configuration. The alternative hypothesis was that they come from a distribution with a different variance.

## 3 Results

Twenty young, normal-hearing participants (17 females and 3 males; mean age 24.3 years) were included in the study (see Borges et al., 2024 [5] for details). In a joint analysis, including all participants and each electrode configuration examined, SNR was a significant predictor of reconstruction accuracy (for details on the statistical test and linear mixed model results, see Supplementary Material S1)

The results from the SRT_neuro_ estimation are shown in figure 4 and table 1, for all accepted sigmoid fits see Supplementary Material S2. The assessment of a relative increase in estimation quality was based on several factors: an increased number of participants for whom it was possible to estimate the SRT_neuro_, an increased number of participants with a SRT_neuro_ within 3 dB from the SRT_beh_, and a decreased SD of the difference between the SRT_neuro_ and the SRT_beh_. A non-significant difference of variance between an in-ear and/or around-the-ear electrode configuration and the Scalp configuration indicates good estimation quality, as the Scalp configuration with its greater spatial coverage was expected to yield the most precise SRT_neuro_ estimate and thus served as a reference.

**Table 1:** Comparison between the EEG-based SRT_neuro_ and the behavioral SRT_beh_ for the examined electrode configurations. The first row indicates the number of participants for whom the SRT measure was successfully estimated from EEG. In the following rows, the numbers of participants for whom the SRT_neuro_ was within 3 dB from the SRT_beh_ are shown along with the median difference between the SRT_neuro_ and the SRT_beh_, the standard deviation of that difference, and the p-value obtained from chi-square test comparing the variance of the difference between the SRT_beh_ and the SRT_neuro_ for each of the in-and-around-the-ear electrode configurations with the corresponding variance obtained for the Scalp configuration..

|  | Ear | AEar | M | T | Earm | AEarm | Eart | AEart | MT | EarmT | AEarmT | LAearMT | RAearMT | Scalp |
| --- | --- | --- | --- | --- | --- | --- | --- | --- | --- | --- | --- | --- | --- | --- |
| **No of estimations** | 9 | 8 | 7 | 14 | 14 | 11 | 19 | 17 | 18 | 20 | 20 | 13 | 19 | 20 |
| **<3 [dB]** | 6 | 6 | 5 | 11 | 6 | 7 | 11 | 14 | 17 | 14 | 19 | 10 | 19 | 19 |
| **Median [dB]** | -1.55 | -0.44 | -0.05 | 0.22 | -1.37 | 0.09 | 0.10 | -0.08 | 0.48 | -1.49 | -0.28 | -1.26 | 0.15 | -0.16 |
| **SD [dB]** | 3.05 | 2.58 | 2.83 | 2.58 | 6.80 | 4.21 | 6.75 | 2.04 | 1.65 | 2.76 | 1.80 | 4.58 | 1.56 | 1.37 |
| **p-value** | $<10^{-3}$ | 0.002 | $<10^{-3}$ | $<10^{-3}$ | $<10^{-3}$ | $<10^{-3}$ | $<10^{-3}$ | 0.007 | 0.200 | $<10^{-3}$ | 0.049 | $<10^{-3}$ | 0.336 | - |

**Figure 4.**
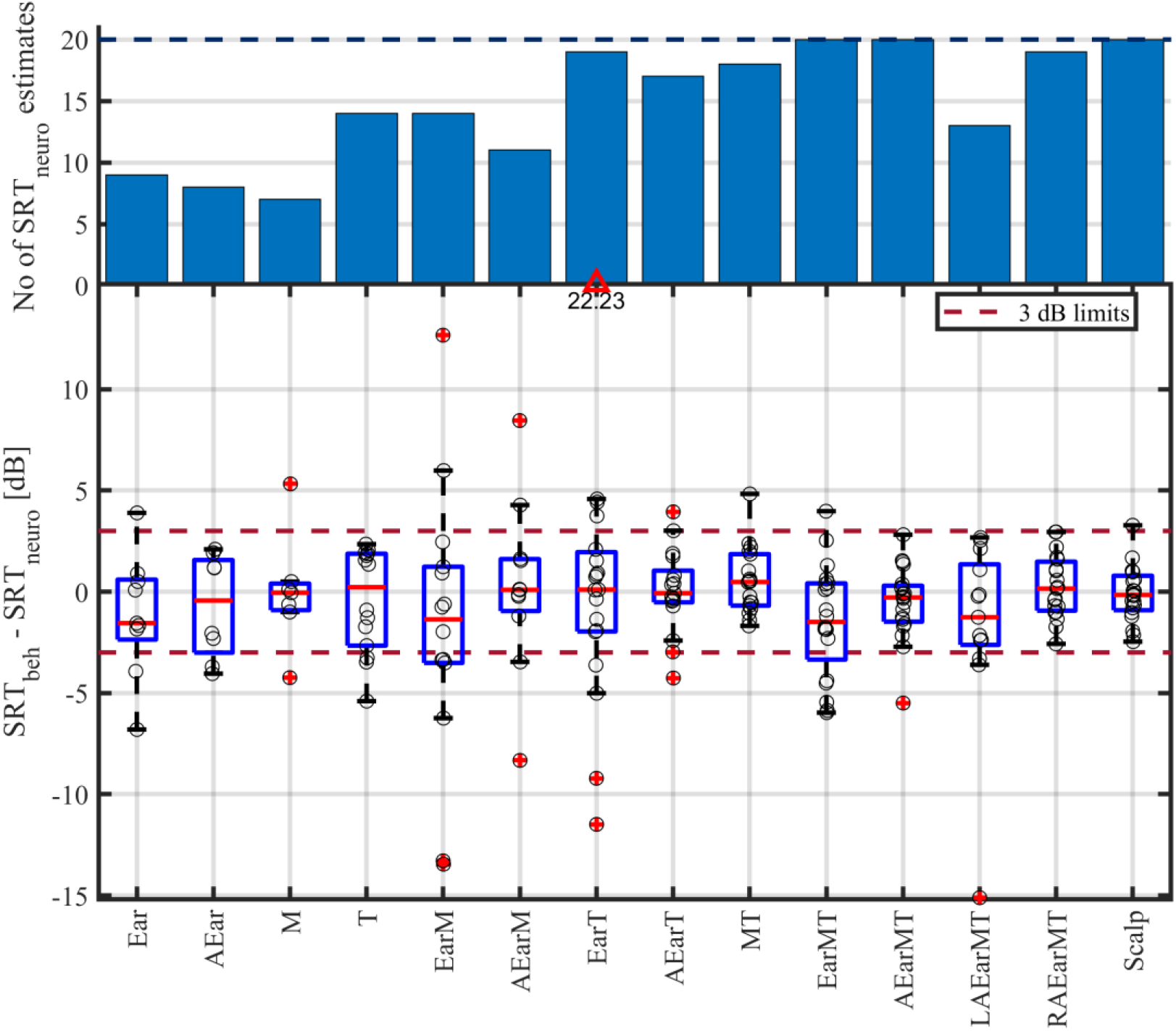
Upper panel shows a bar plot depicting the number of participants for whom a valid SRT_neuro_ were obtained. The blue dotted line indicates the number of participants included in the study. The lower panel illustrates the difference between the (estimated) SRT_neuro_ and the (measured) SRT_beh_ for the considered electrode configurations. The difference in dB for each participant is marked with black circles for each of the electrode configurations, with a slight horizontal jitter. A boxplot of the difference is also shown, where the red line indicates the median, the edge of the blue box indicates the 25th and 75th percentiles and the black whiskers mark the most extreme values for non-outliers. Datapoints are considered outliers if they are more than 1.5 times the interquartile range away from the bottom or top of the box and marked with a red cross; extreme outliers outside the depicted value range are indicated with a red triangle.

For the Scalp configuration a SRT_neuro_ was found for all 20 participants (100%) and the SRT_neuro_ was within 3 dB difference of the behavioral SRT_beh_ for 19 participants (95%; see figure 4 and table 1). The SD of the difference was 1.37 dB. In contrast, using only ear-EEG electrodes (Ear and AEar) resulted in 9 (45%) and 8 (40%) SRT_neuro_ estimates, respectively, with only 6 participants (30%) showing a difference within 3 dB. The Ear configuration yielded an SD of the difference of 3.05 dB and the AEar configuration an SD of the difference of 2.58 dB. When using only the mastoids as input to the decoder (M), the results were similar to those of the Ear and AEar configurations, with 7 (35%) estimates and 5 (25%) participants showing a difference within 3 dB and an SD of 2.83 dB. However, using temporal electrodes alone (T) demonstrated improved predictive power compared to the M, Ear and AEar configuration, as indicated by the higher number of participants for whom a SRT_neuro_ could be estimated (14/70%) and the higher number of participants with the SRT_neuro_ within a 3 dB of the SRT_beh_ (11/55%). The SD for the T configuration was identical to the AEar configuration (but based on more participants) and smaller than that obtained for the Ear and M configuration.

When combining the electrodes in the AEar or Ear configuration with the mastoids (AEarM and EarM), the SRT_neuro_ estimates increased compared to using (averaged) ear electrodes or mastoids alone (AEar, Ear and M). However, the number of participants within 3 dB showed only minor differences, and the SD increased, indicating that the additional SRT_neuro_ estimates were not very precise. By contrast, combining the temporal electrodes with the averaged in-ear electrodes (AEarT) or non-averaged in-ear electrodes (EarT) resulted in an overall improvement for all parameters compared to using AEar, Ear or T separately. The only exception was the SD for the EarT configuration, which increased compared to using T or Ear alone and due to an extreme outlier. Additionally, when combining the temporal electrodes with the mastoids (MT), there was an improvement in all parameters compared to using the electrodes individually (M and T).

The electrode configurations combining the in-ear electrodes, mastoids, and temporal electrodes (EarMT, AEarMT) yielded some improvement in predictive power compared to the configurations containing a combination of Ear and T or Ear and M electrodes (AEarM, AEarT, EarM, EarT). The improvement provided by AEarMT (relative to AEarM and AEarT) far exceeded that provided by EarMT (relative to EarM and EarT). The estimated SRT_neuro_ within 3 dB of the SRT_beh_ was 19 (95%) for the AEarMT configuration. When comparing EarMT and AEarMT with MT, AEarMT exhibited better predictive power regarding participants within 3 dB and number of estimates, although the standard deviation increased slightly compared to the MT configuration. Conversely, the EarMT configuration showed reduced predictive power compared to MT across all parameters except for the number of possible estimates. Overall, the performance obtained with the AEarMT configuration was largely on par with that obtained using the reference Scalp configuration.

When focusing only on electrodes from one hemisphere (LAEarMT and RAEarMT), it was found that the information from the right hemisphere (RAEarMT) was sufficient to match the performance obtained when using information from both hemispheres (AEarMT). However, relying solely on information from the left hemisphere (LAEarMT) led to a reduction in estimation accuracy relative to AEarMT across all considered measures, as the amount of participants within 3 dB of the SRT_beh_ decreased to 10 (50%), the SD increased substantially, and the variance of the difference was significantly larger compared to the Scalp configuration. When analyzing the variance of the differences between the SRT_neuro_ and the SRT_beh_, no significant difference (α =0.05) was found between the Scalp configuration and the configurations MT and RAEarMT whereas the otherwise very competitive AEarMT configuration yielded an SD that was just significantly different than that obtained from the Scalp configuration.

## 4 Discussion

Overall, the number of successful SRT estimations increased with the inclusion of electrodes from more locations (see figure 4 and table 1), as did the number of participants for whom the SRT_neuro_ fell within 3 dB of the SRT_beh_. This increase was particularly pronounced with addition of the temporal electrodes (T). Averaging the in-ear electrodes (AEar) generally increased the predictive power of the SRT_neuro_, increasing the number of participants within 3 dB difference from the SRT_beh_ and reducing the standard deviation (SD) of the difference. Additionally, the median of the difference remained within 2 dB for all electrode configurations and was generally very close to zero.

On the group level, the SNR was a significant predictor of reconstruction accuracy across all electrode configurations in the study, although it was not possible to determine the SRT_neuro_ value for all participants in every configuration. This may be attributed to individual differences in brain anatomy and the quality of the recorded signal [23] as well as differences in physiological responses. In a previous study by Borges et al. 2024 [5], the SRT_neuro_ values for scalp EEG were estimated using the same data as used in the current study, obtained with 66 scalp electrodes. The results closely matched the Scalp results stated here, with all participants showing a difference within 3 dB, with an SD of 1.45 dB, and a median of 0.4 dB. The slight difference compared to the “Scalp” electrode configuration reported here stems from the different EEG preprocessing pipelines used: the previous study used an independent-component-analysis (ICA) based artefact rejection method [24] and an average reference, which were not used in the current study. The ICA-based artefact rejection method was not included in this study to ensure that both the in-ear-EEG and the scalp EEG underwent the same preprocessing steps. Since the ICA-based artefact rejection relies on the spatial distribution of the individual components and is trained on whole-scalp EEG topographies [24], it is not suitable for ear-EEG which only covers a narrow part of the scalp. In Borges et al. (2024) [5], the sigmoid fit was only performed if the reconstruction accuracy for the clean speech condition was significantly higher than the noise floor. In the current study, the sigmoid fit was only performed if the reconstruction accuracy of the high-SNR conditions (clean speech, SRT_beh_ +2dB, and SRT_beh_ +4dB) was higher than that of the low-SNR conditions (SRT_beh_ -2dB, SRT_beh_ -4dB and the noise floor). This adjustment was necessary because using fewer electrodes in different configurations introduced more noise, requiring additional data to confirm an increase in reconstruction accuracy with SNR. Due to the supplementary noise in some of the electrode configurations additional measures to ensure a SRT_neuro_ of good quality were implemented, namely rejection of unlikely SRT_neuro_ values (outside the range [-40, 40] dB) and rejection fits were the average reconstruction accuracy for a condition was above the average clean speech reconstruction accuracy for more than two SNR conditions or below the noise floor average reconstruction accuracy for more than two conditions.

Prediction of the SRT_neuro_ values was also conducted in a study by Lesenfants et al. (2019)[4], where they obtained a SRT_neuro_ within 2 dB from the SRT_beh_ for 8 out of 19 participants (47%) using a delta-band envelope-based encoder. In the current study 17 out of 20 participants (85%) obtained a SRT_neuro_ within 2 dB from the SRT_beh_ when using the Scalp configuration. Both the speech material (used for behavioral testing and EEG stimuli) and the model used show critical differences to the current study, which may explain the differences in results, for detailed information see the discussion of Borges et al. 2024 [5].

The SRT_neuro_ estimation performance using electrode configurations solely with in-ear electrodes (Ear, AEar) was substantially lower than that of the Scalp configuration. This reduction could be attributed to several factors, including the fact that the in-ear electrodes are dry electrodes and therefore have a higher impedance compared to the wet electrodes used on the scalp. More likely, the reduction is mostly due to the major reduction of spatial coverage, as the Scalp configuration has a comprehensive coverage of the entire scalp while the in-ear electrodes are limited to inside the ear. However, it was also found that a reduction of spatial information can result in similar results as when using the Scalp configuration if the electrodes are placed in the optimal locations, such as is the case for the AEarMT and RAEarMT configurations (using also around-the-ear electrodes). Averaging the ear electrodes (AEar) systematically resulted in an improved SRT estimation compared to using a non-averaged ear configuration (Ear). This effect is not as obvious when using only in-ear electrodes (Ear, AEar), but becomes evident when combined with scalp electrodes such as AEarMT and EarMT. This improvement could be due to the additional spatial coverage provided by more electrodes inside the ear being less beneficial than averaging the in-ear-electrodes – because averaging the in-ear electrodes would mean achieving a higher SNR and reducing the number of parameters the decoder needs to estimate. This points towards the decoder being unable to provide optimal weighting of the electrodes with the data available in the current study.

Using mastoids only (M) yielded results similar to those obtained with only in-ear electrodes (Ear and AEar). In a previous study by Borges et al. 2024 [5], the encoding analyses of the same stimuli showed a higher prediction accuracy with electrodes in the frontocentral scalp region. This suggests that the mastoid and in-ear electrodes are aligned along approximately the same equipotential line, resulting in comparable signal characteristics. Conversely, using only the temporal electrodes (T) nearly doubled the number of participants with a SRT_neuro_ within the 3-dB difference from the SRT_beh_ as compared to the AEar and Ear configurations, without increasing the SD of the difference between the SRT_neuro_ and the SRT_beh_. This improvement could be attributed to the temporal electrode’s closer proximity to the frontocentral plane compared to the mastoids, as well as their placement directly above the auditory cortex, leading to a better SNR. Additionally, combining averaged in-ear-EEG electrodes with temporal electrodes (AEarT) improved the estimation accuracy compared to AEar and T alone, likely due to better spatial coverage and therefore higher SNR. The improvement observed when combining averaged in-ear electrodes with the temporal electrodes was greater than when combining the averaged in-ear electrodes with the mastoids. This could be attributed to a greater electrical potential difference between the ear and temporal electrodes compared to that between the ear and mastoid, resulting in an enhanced SNR.

A general improvement in estimation quality was observed when combining in-ear-EEG, temporal and mastoid electrodes compared to configurations that omitted any of the electrode groups, except for EarMT, which exhibited a lower estimation quality than MT. The highest estimation quality for configurations with electrodes in and around the ear was found for the AEarMT configuration, closely matching the estimation quality obtained from all scalp electrodes. We further assessed the estimation quality using electrodes from only one hemisphere within the AEarMT electrode configuration, as it represented the best-performing configuration for electrodes positioned in and around the ear. When using the right-side electrodes only (RAEarMT), the performance remained largely the same as obtained with AEarMT using both hemispheres, as the number of estimates within a 3 dB difference remained the same, the SD decreased, and the difference in variance compared to the Scalp configuration was non-significant, which was not the case for the AEarMT configuration (p=0.049). This suggests that RAEarMT and AEarMT yield comparable results. Conversely, using only left-sided electrodes (LAEarMT) led to a drop in the number of the SRT_neuro_ within 3 dB, alongside a large increase in SD, indicating a worse SRT prediction performance. Borges et al. (2024)[5] already showed that the average prediction accuracy is more pronounced in the right hemisphere (using the same data set as used in the present study), potentially resulting in a higher SNR and a better estimation quality. This is further supported by findings from Abrams et al. 2008 [25], which demonstrated a strong right hemisphere dominance in speech envelope coding.

### Application

The SRT can be estimated using in-ear EEG alone, and the findings of this study suggest that accurate SRT estimation is feasible with electrodes positioned in and around one ear, showing similar performance when using electrodes exclusively from the right side. This means that if incorporating the SRT_neuro_ estimation in hearing aids, they do not need to be electrically connected by a wire. While for the configurations using only in-ear EEG electrodes this is currently true only for a subset of participants. Further improvement of the method — such as training the model on more data to enhance SNR [23,26] adding more predictors such as phoneme onset and spectrogram [4], or utilizing more complex nonlinear decoding models [27] — could enable a SRT_neuro_ integration in hearing aids. Assuming that the SRT estimation also works with non-controlled stimuli, the SRT could be constantly assessed and utilized as a real-time control signal in the hearing aids, allowing, e.g., noise-reduction algorithms to be adjusted dynamically to lower the SRT when needed for the individual user. Additionally, continuous SRT logging in hearing aid users’ natural environments could further support hearing rehabilitation. Estimating the SRT in real-world conditions would likely require more data - and thus more time - than in a controlled clinical setting. However, this may not be a limitation for real-life applications, where significantly more time is available than in a typical clinical assessment. The potential benefits of adding electrodes to hearing aids are also supported by the ability to decode the attended speaker [11,12,17,28,29], which would allow for user controlled beamformer adaption.

### Limitations

There are several limitations to the SRT_neuro_ estimation method. First, while the envelope following response is closely linked to speech intelligibility [3,30–32], it is not a direct measure of speech intelligibility itself. This means that using speech material in a language unfamiliar to the participant an SRT_neuro_ estimate could potentially still be conducted, making the estimated SRT_neuro_ more of an evaluation of access to the speech envelope information than an accurate measure of speech intelligibility itself. Second, the study is limited by the amount of data recorded. Since ear-EEG is suitable for long-term recordings [6], collecting more data could improve the SNR in the in-ear EEG measurement [23] potentially enhancing the SRT_neuro_ estimation precision beyond what was observed in the current study. Third, if the SRT_neuro_ is to be estimated in the hearing-aid users’ natural environment, additional factors such as different noise types, target speech and different populations must be considered to ensure accurate estimations.

Since people with hearing impairment is the target group of the application of measuring SRT_neuro_ and the most common type of hearing impairment is age related [33], it is important also to investigate the effect of age and hearing status on the SRT_neuro_ method. Previous studies [34,35] have shown that there is an increase of reconstruction accuracy with age. While the sigmoid fitting method used in the current study to find the SRT_neuro_ is independent of the absolute reconstruction accuracy values, it is interesting to see whether there are any changes in the precision of SRT_neuro_ method. Another study by Presacco et al. (2019) [36] compared the envelope following response of an old hearing impaired and old normal hearing population, here they did not find an overall difference of envelope reconstruction accuracy between the two groups. However, testing the methods used in the current study on a hearing-impaired population would provide SRT_beh_ with more variance and therefore further test the robustness of the SRT_neuro_ method used.

## 5 Conclusion

The SRT values were successfully estimated from ear-EEG in a subgroup of the tested population. Additionally, the SRT estimation accuracy improved when combining in-ear electrodes with electrodes around the ear. Notably, using electrodes from only the right side of the head yielded performance similar to that achieved with electrodes from both sides. Future work should investigate the effect of age and hearing impairment on the SRT_neuro_ estimation.

## Data availability statement

The data used in the study is available upon reasonable request to the corresponding author.

## Ethical statement

The study was conducted in accordance with the principles embodied in the Declaration of Helsinki and in accordance with local statutory requirements. It was approved by the Institutional Review Board at Aarhus university with the approval number TECH-2022-004. Written informed consent was obtained for all participants prior to participation in the study. The participants were informed that the results would be published in scientific journals, that they could leave the study at any time without providing a reason, and that they can request their data removed prior to anonymization.

## Conflict of interest

The authors have no conflicts of interest to declare.

## Data Accessibility

Data can be provided upon reasonable request.

## Acknowledgement

The authors are very thankful for the participants who participated in the study and for Simon With’s great support during data collection. Lastly, they are thankful for the funding from the William Demant Foundation [Grant number 21-2912].

## CrediT authorship contribution statement

Heidi B. Borges

Writing – original draft, Data curation, Formal analysis, Funding acquisition, Conceptualization, Software, Investigation, Visualization, Methodology. Johannes Zaar: Writing – review and editing, Funding acquisition, Conceptualization, Supervision, Methodology, Software, Project administration. Emina Alickovic: Writing – review and editing, Funding acquisition, Conceptualization, Supervision, Methodology, Software. Christian B. Christensen: Writing – review and editing, Conceptualization, Supervision, Methodology, Resources. Preben Kidmose: Conceptualization, Methodology, Writing – review and editing, Supervision, Project administration, Funding acquisition.

## Notes

### Competing Interest Statement

The authors have declared no competing interest.

