## Supplementary material for "The Speech Reception Threshold Can be Estimated Using EEG Electrodes In and Around the Ear"

### S1: The results from the linear mixed-effects model

| Configuration | Estimate of SNR coefficient | Standard error | *t*-value | *p*-value |
|---|---|---|---|---|
| **Ear** | 0.0015 | 0.0004 | 4.145 | $<10^{-3}$ |
| **AEar** | 0.0018 | 0.0004 | 4.770 | $<10^{-3}$ |
| **M** | 0.0023 | 0.0004 | 6.383 | $<10^{-3}$ |
| **T** | 0.0028 | 0.0004 | 6.816 | $<10^{-3}$ |
| **EarT** | 0.0030 | 0.0004 | 7.518 | $<10^{-3}$ |
| **AEarT** | 0.0042 | 0.0004 | 10.132 | $<10^{-3}$ |
| **MT** | 0.0048 | 0.0004 | 11.175 | $<10^{-3}$ |
| **EarM** | 0.0017 | 0.0004 | 4.706 | $<10^{-3}$ |
| **AEarM** | 0.0023 | 0.0004 | 5.813 | $<10^{-3}$ |
| **EarMT** | 0.0035 | 0.0004 | 9.445 | $<10^{-3}$ |
| **LAEarMT** | 0.0024 | 0.0003 | 7.900 | $<10^{-3}$ |
| **RAEarMT** | 0.0038 | 0.0004 | 9.759 | $<10^{-3}$ |
| **AEarMT** | 0.0048 | 0.0004 | 12.062 | $<10^{-3}$ |
| **Scalp** | 0.0094 | 0.0005 | 18.564 | $<10^{-3}$ |

*1: Results from the linear mixed-effects model R ~ SNR + (P|1), where R is reconstruction accuracy, SNR is the signal to noise ratio in the condition and P is a subject specific offset. The results seen are the estimate of the SNR coefficient, the standard error, t-statistics and p-value for each of the electrode configurations from the linear mixed model and Wald test.*

### S2 Accepted sigmoid fits

Underneath the reconstruction accuracies and sigmoid fits are shown for all the accepted sigmoid fits.

Ear

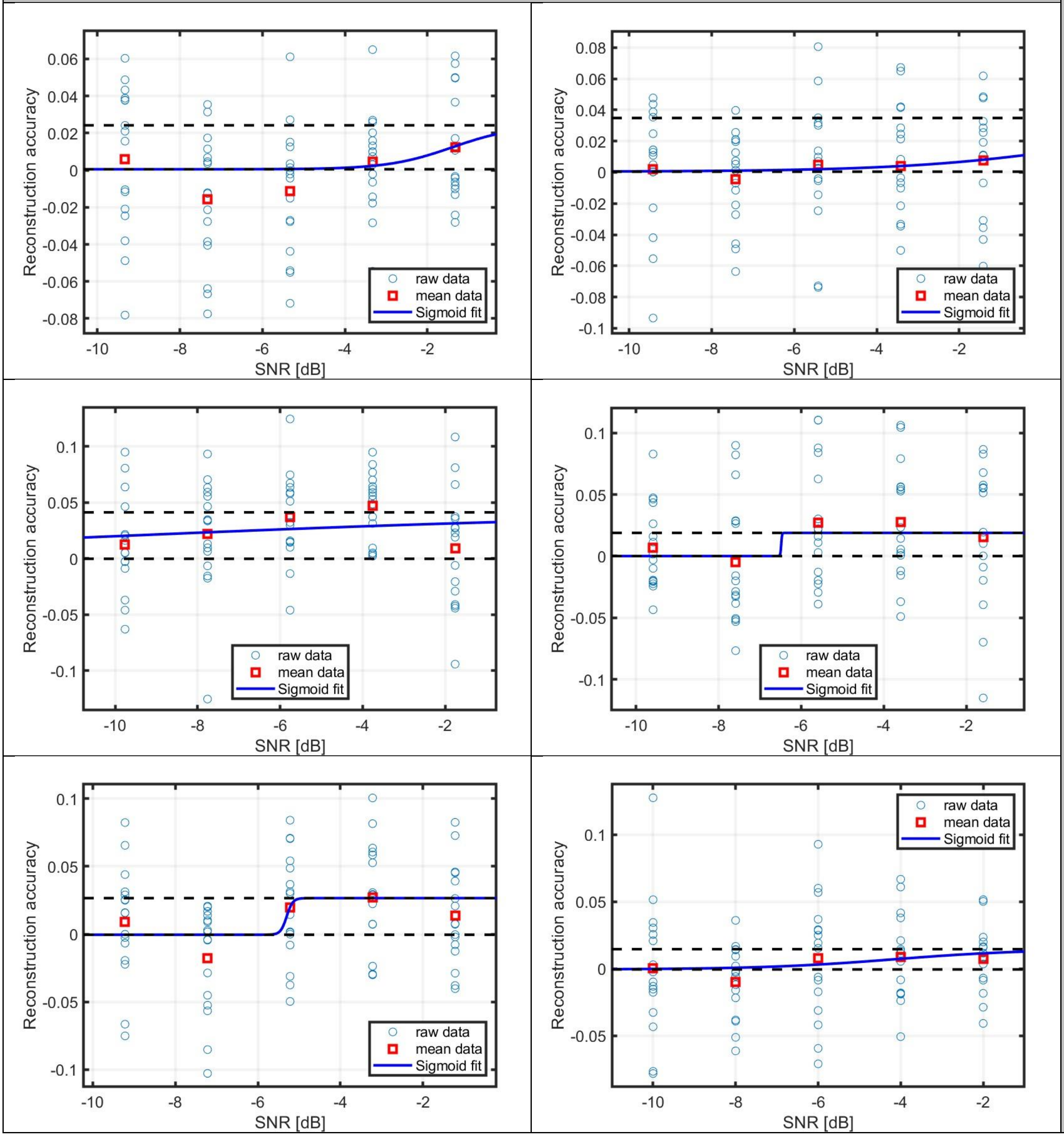


Ear

Ear

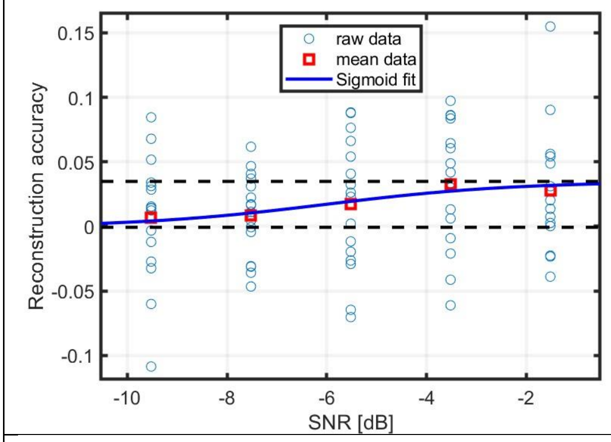


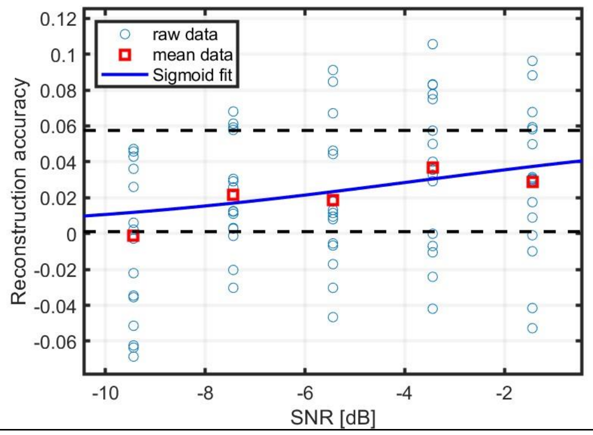


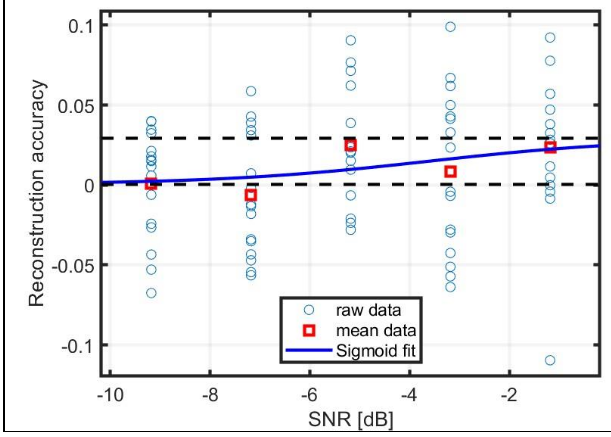


AEar

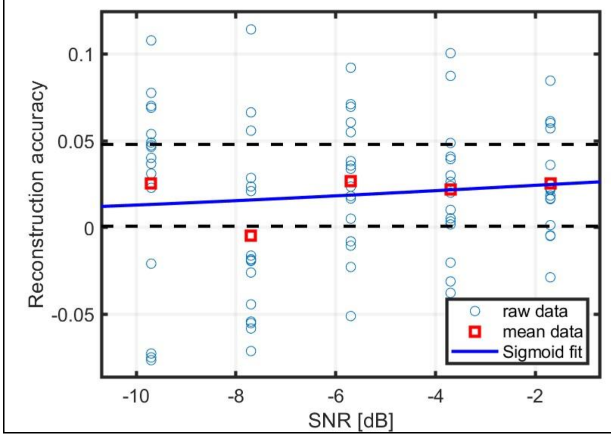


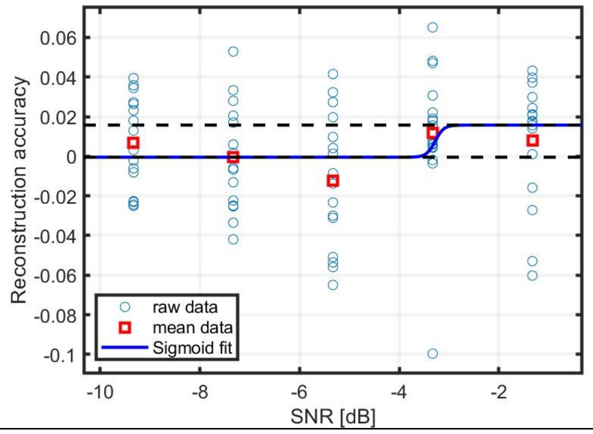


AEar

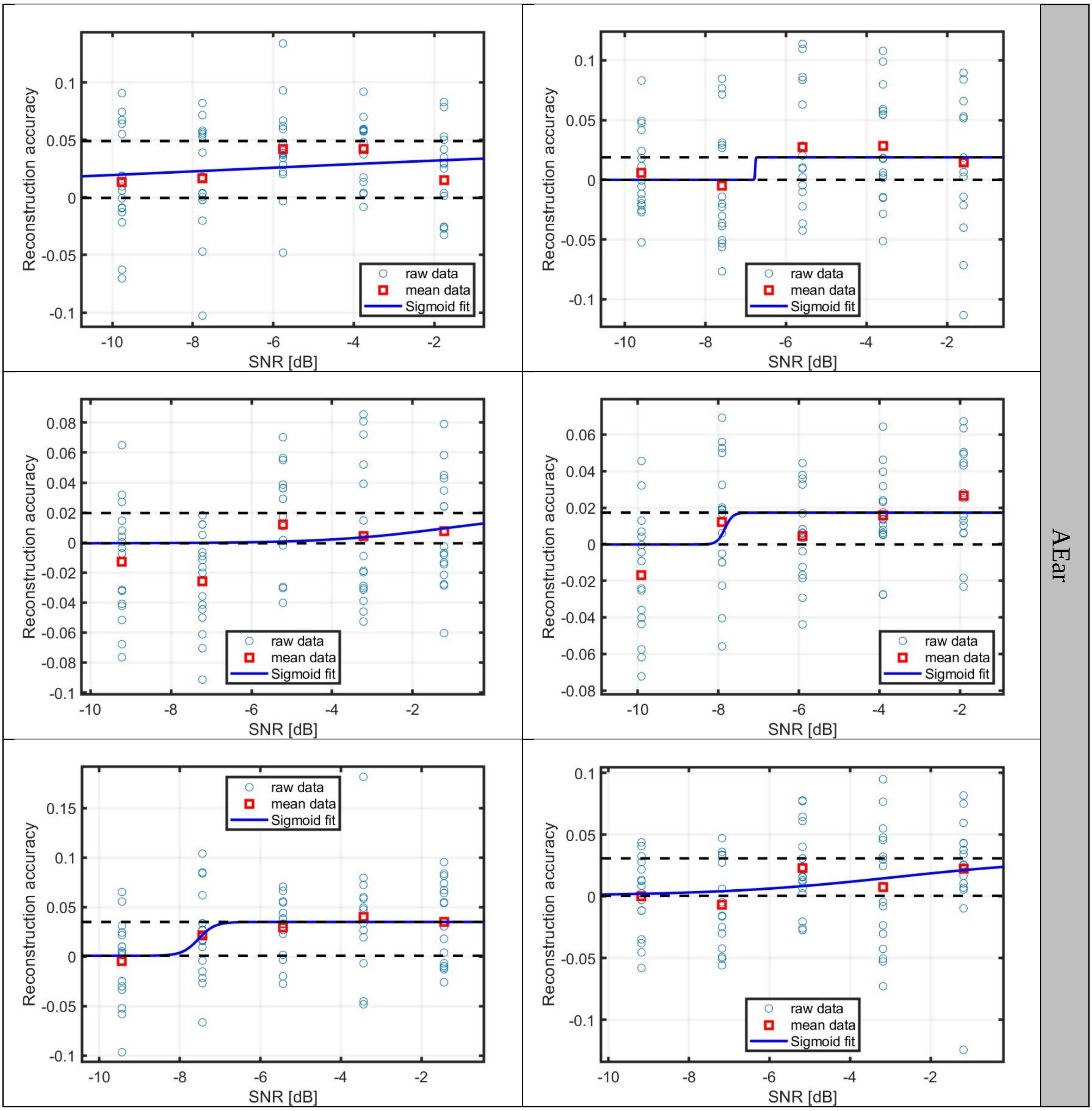
Reconstruction accuracy
SNR [dB]
raw data
mean data
Sigmoid fit
AEar

M

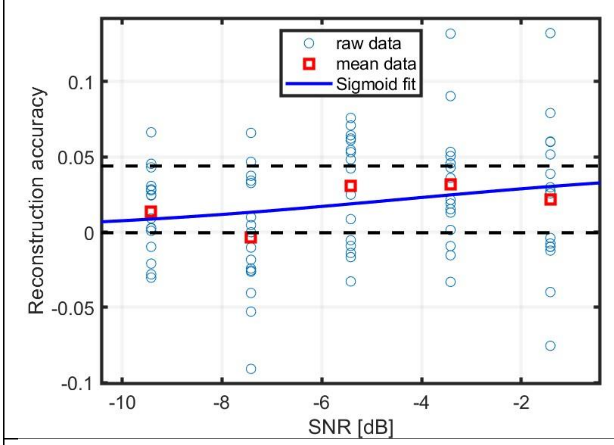
raw data
mean data
Sigmoid fit
Reconstruction accuracy
SNR [dB]

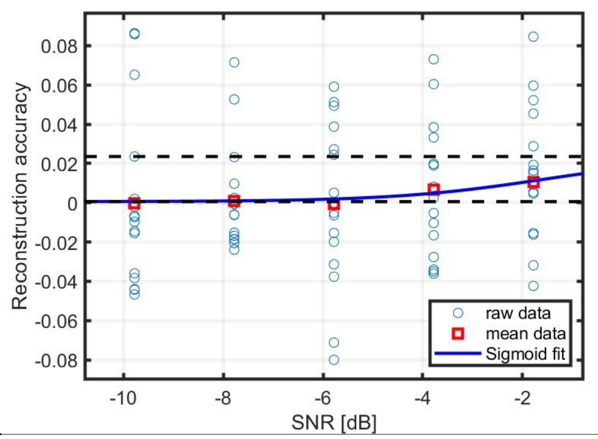
raw data
mean data
Sigmoid fit
Reconstruction accuracy
SNR [dB]

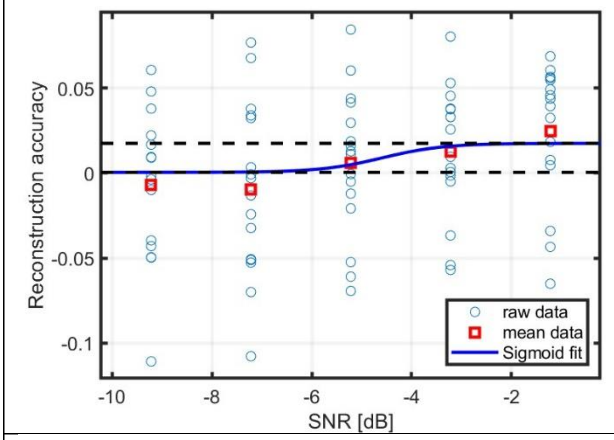
raw data
mean data
Sigmoid fit
Reconstruction accuracy
SNR [dB]

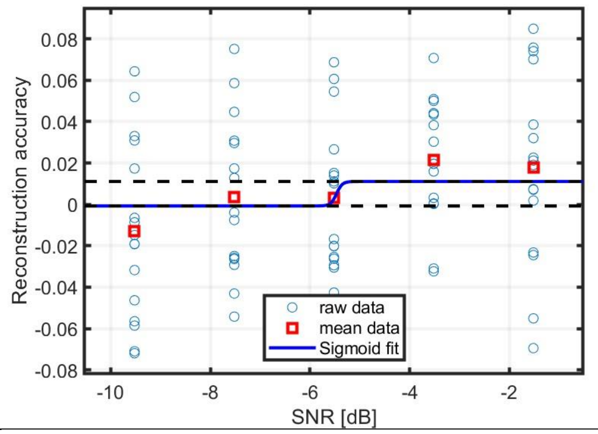
raw data
mean data
Sigmoid fit
Reconstruction accuracy
SNR [dB]

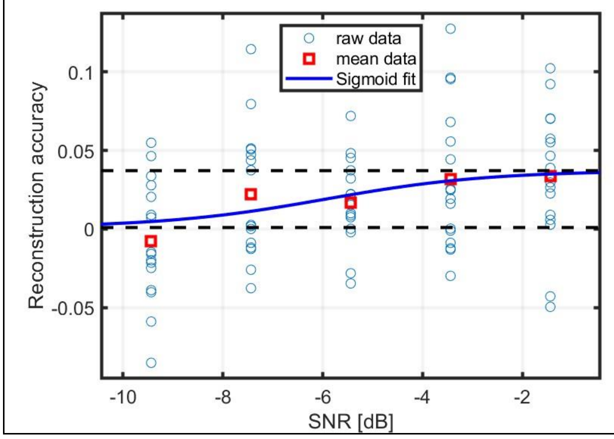
raw data
mean data
Sigmoid fit
Reconstruction accuracy
SNR [dB]

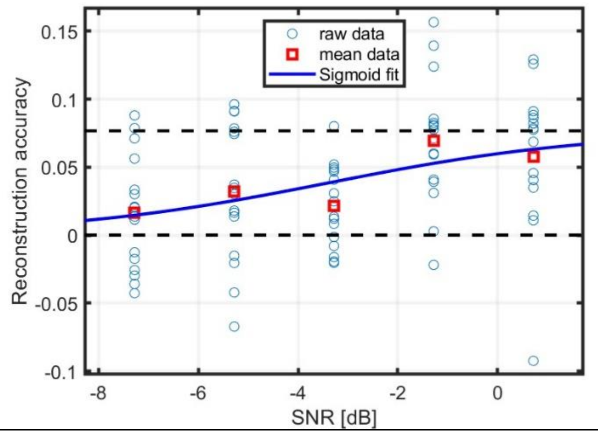
raw data
mean data
Sigmoid fit
Reconstruction accuracy
SNR [dB]

M

<table>
<tr><td>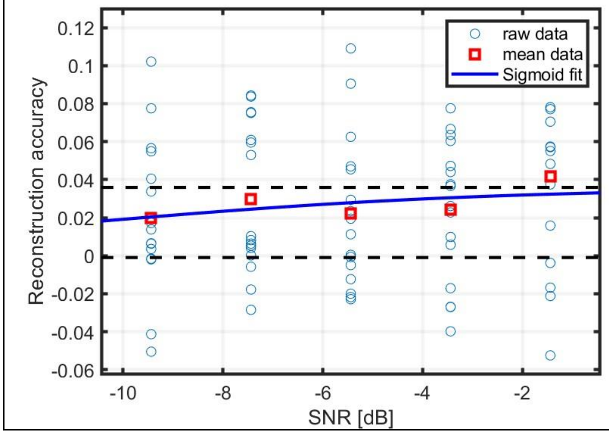
</td><td></td><td>M</td></tr>
</table>

<table>
<tr><td colspan="2">T</td><td rowspan="3">T</td></tr>
<tr><td>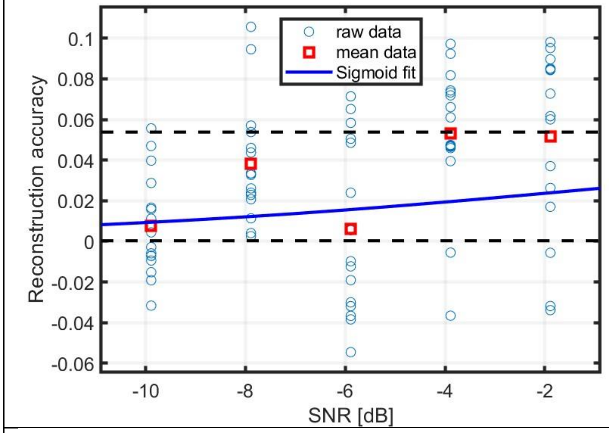
</td><td>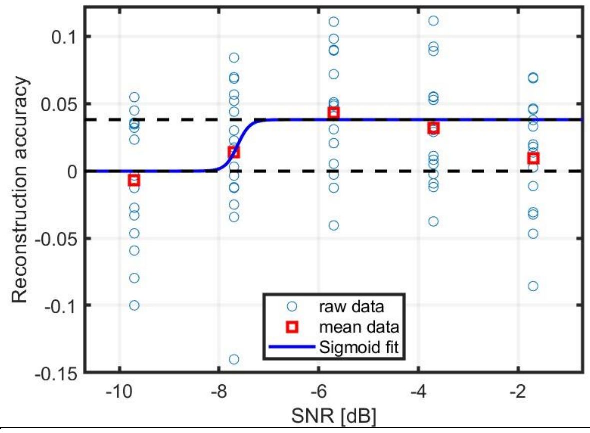
</td></tr>
<tr><td>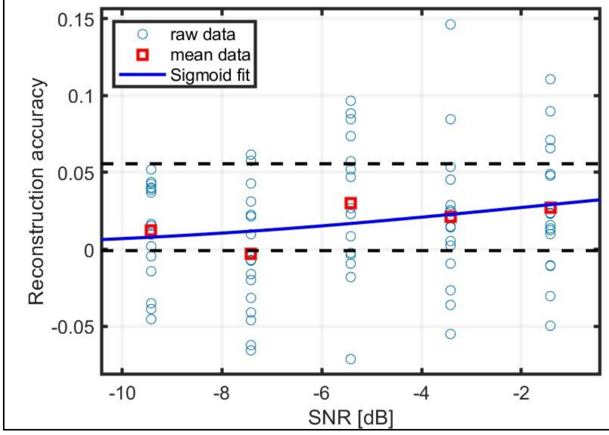
</td><td>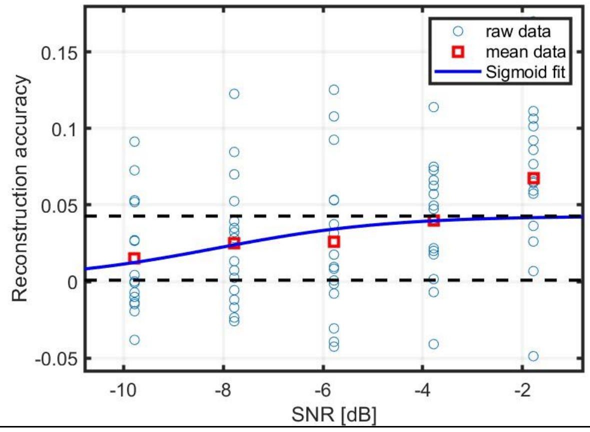
</td></tr>
</table>

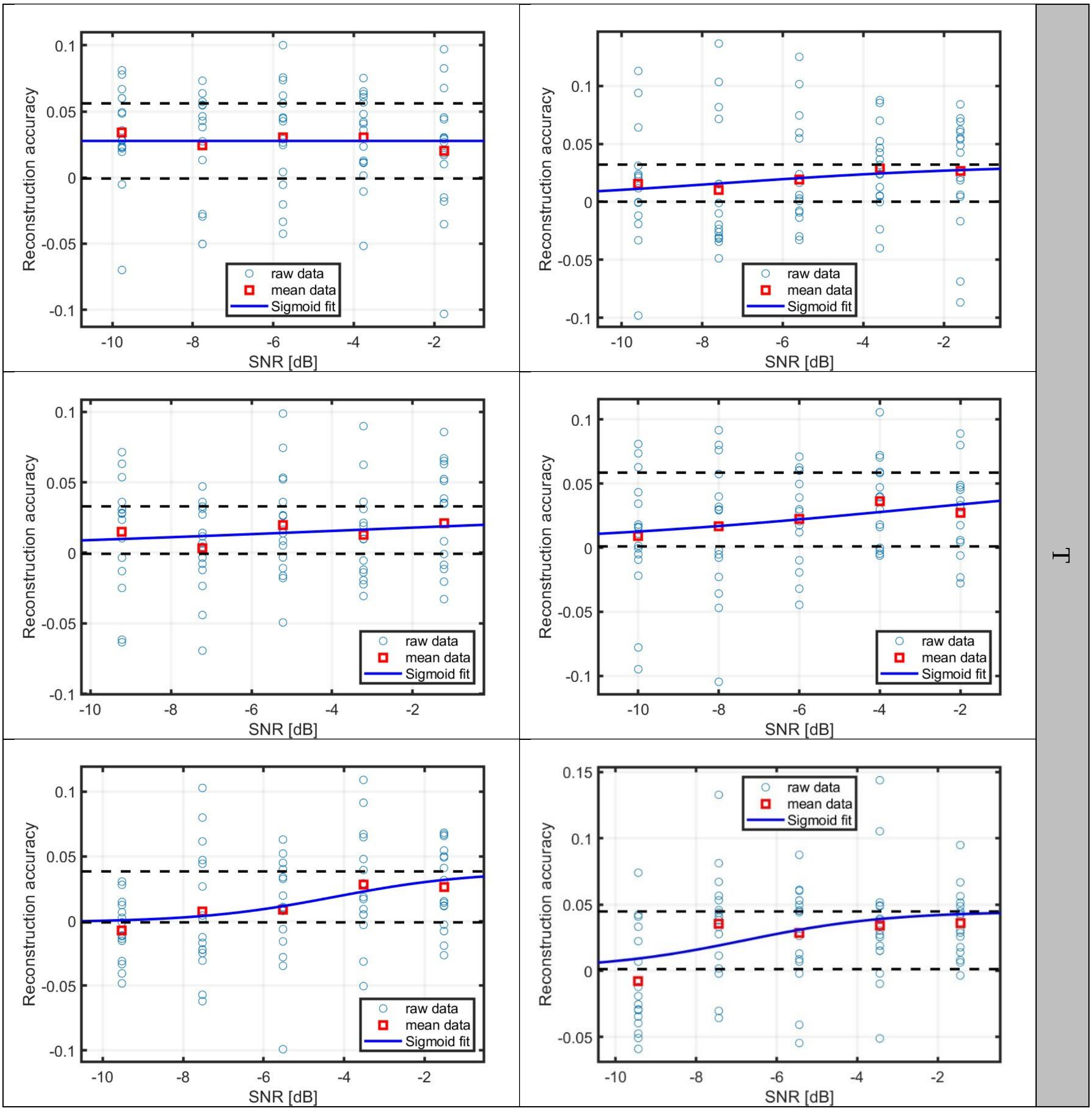
Reconstruction accuracy
SNR [dB]
raw data
mean data
Sigmoid fit
T

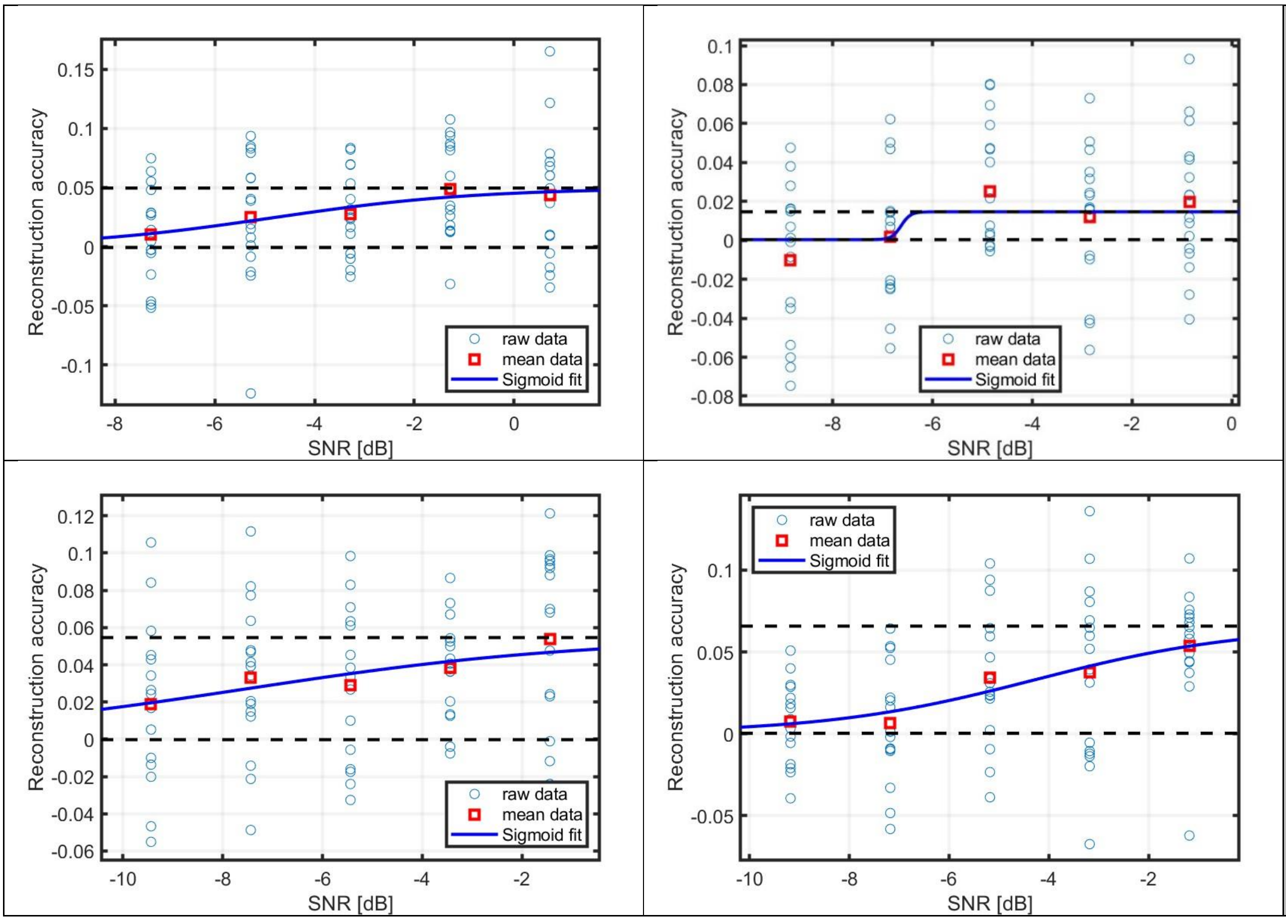
Reconstruction accuracy
SNR [dB]
raw data
mean data
Sigmoid fit

T

EarM

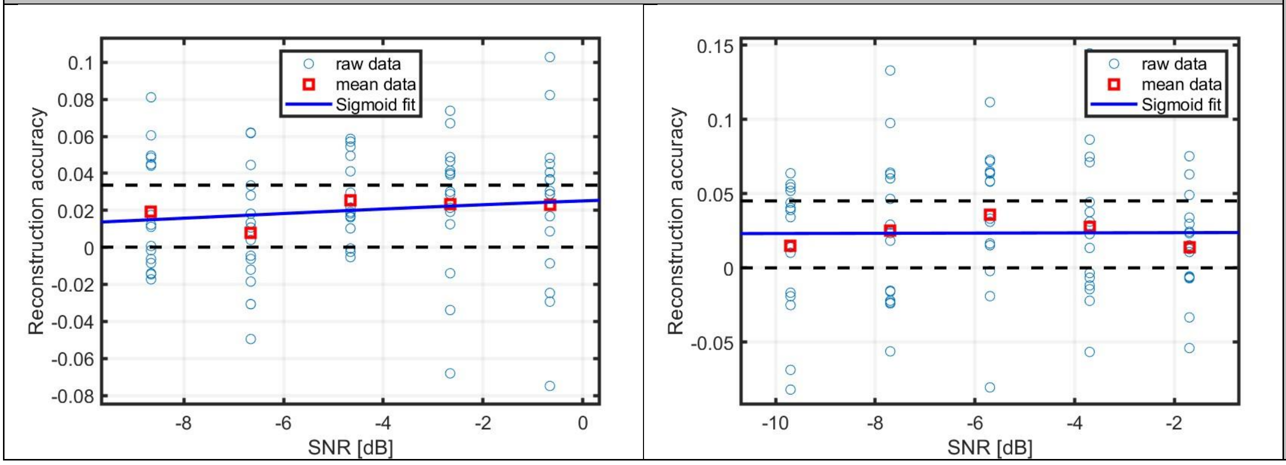
Reconstruction accuracy
SNR [dB]
raw data
mean data
Sigmoid fit

EarM

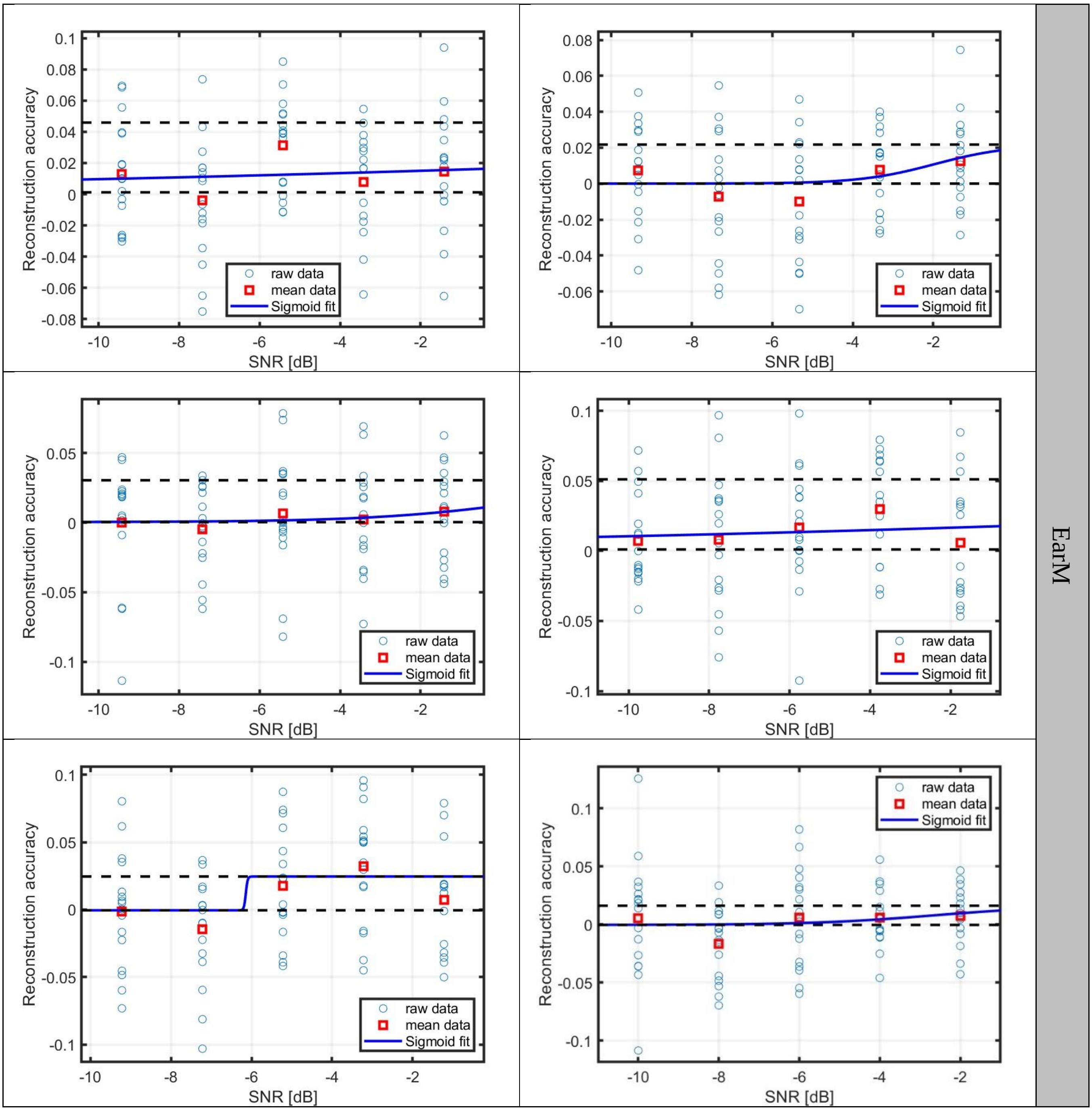
Reconstruction accuracy
SNR [dB]
raw data
mean data
Sigmoid fit
EarM

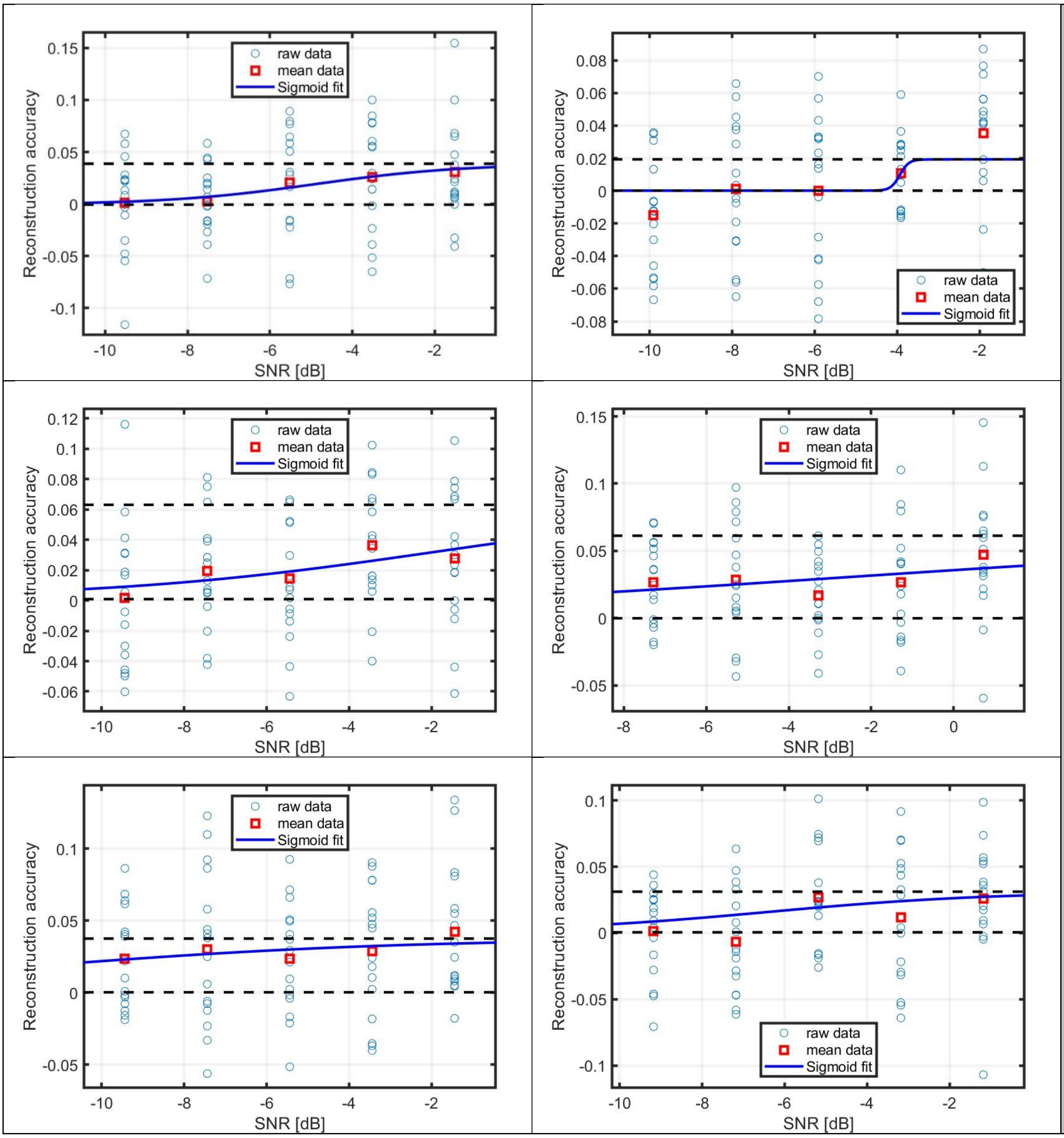
raw data
mean data
Sigmoid fit
Reconstruction accuracy
SNR [dB]

EarM

AEarM

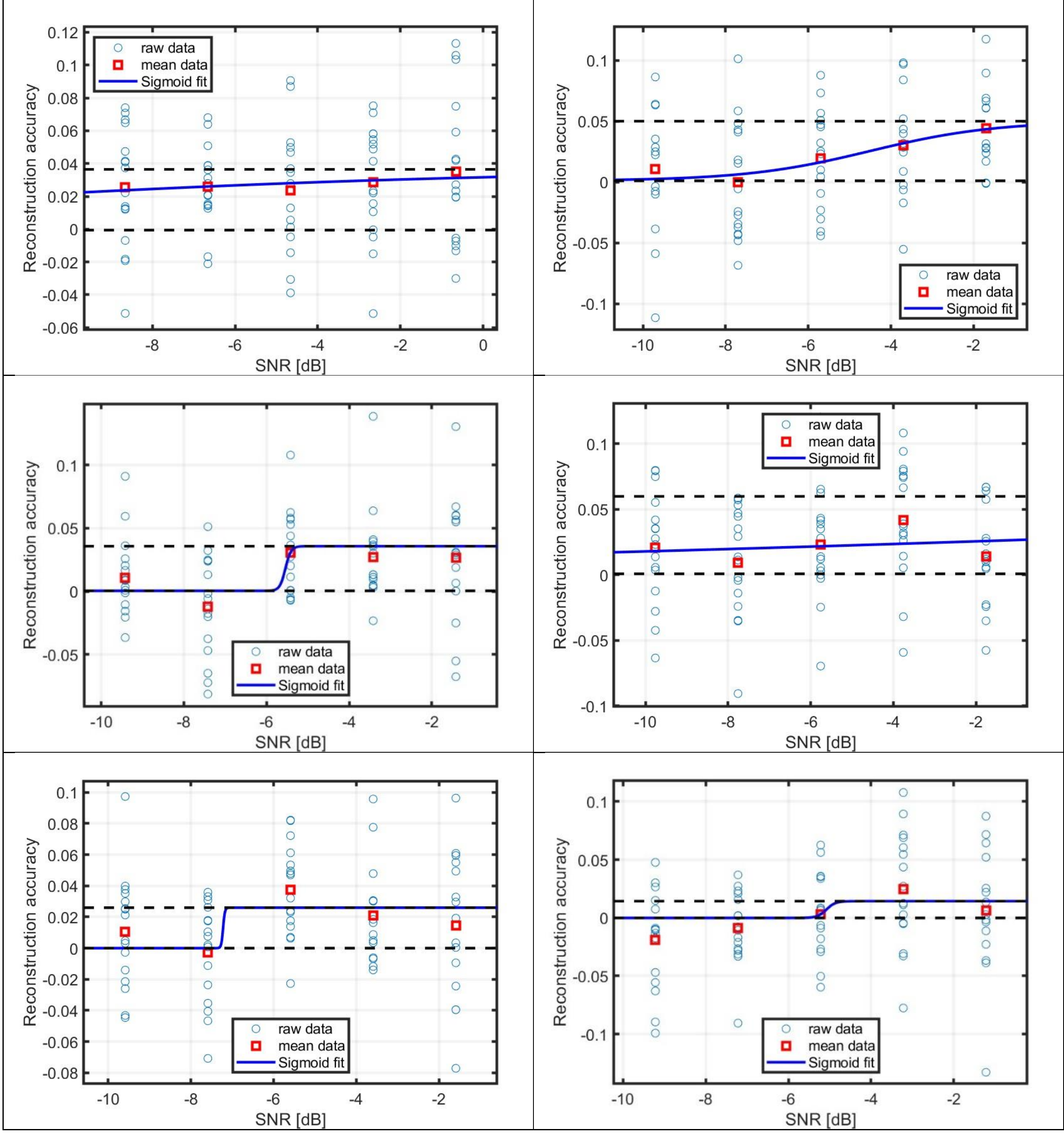
raw data
mean data
Sigmoid fit
Reconstruction accuracy
SNR [dB]
0.12
0.1
0.08
0.06
0.04
0.02
0
-0.02
-0.04
-0.06
-8
-6
-4
-2
0
-0.05
-0.1
0.05
-10

AEarM

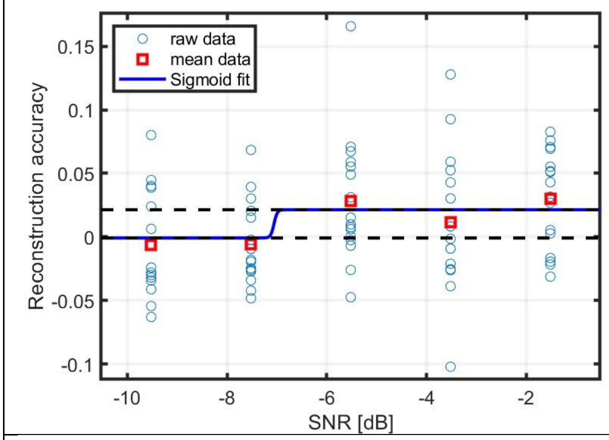
raw data
mean data
Sigmoid fit
Reconstruction accuracy
SNR [dB]

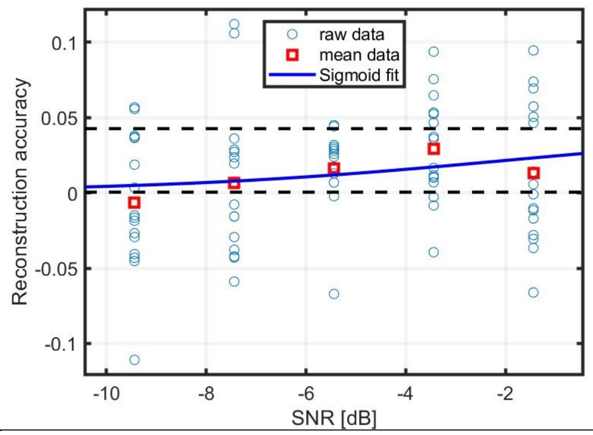
raw data
mean data
Sigmoid fit
Reconstruction accuracy
SNR [dB]

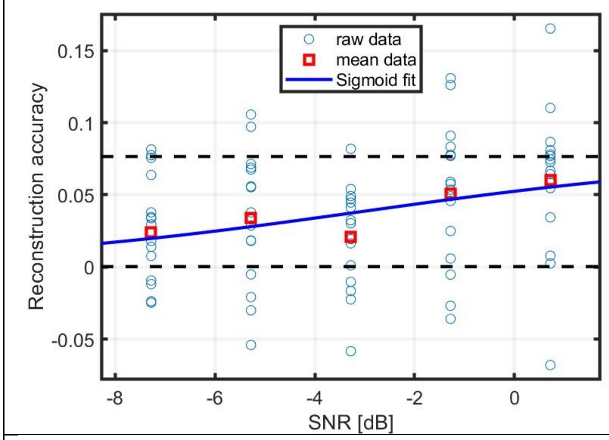
raw data
mean data
Sigmoid fit
Reconstruction accuracy
SNR [dB]

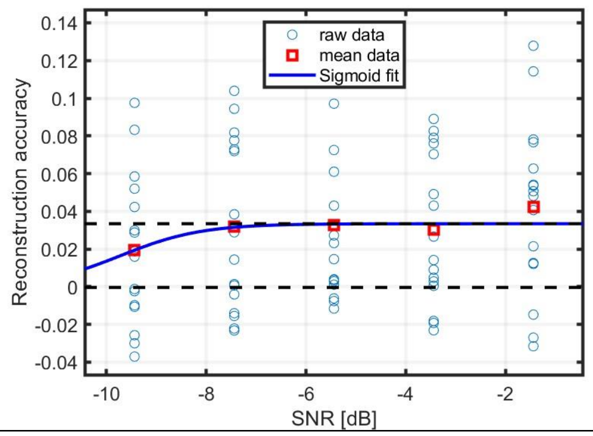
raw data
mean data
Sigmoid fit
Reconstruction accuracy
SNR [dB]

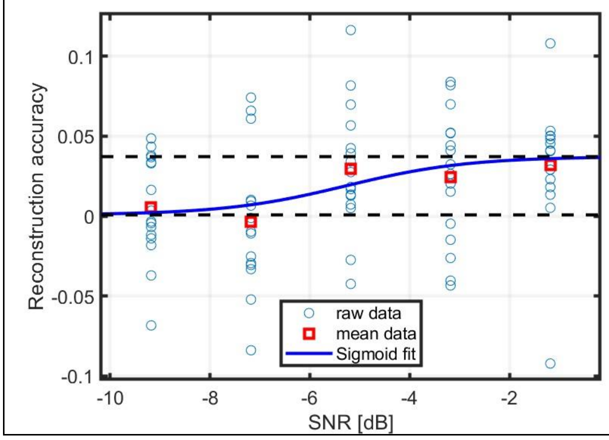
raw data
mean data
Sigmoid fit
Reconstruction accuracy
SNR [dB]

AEarM

EarT

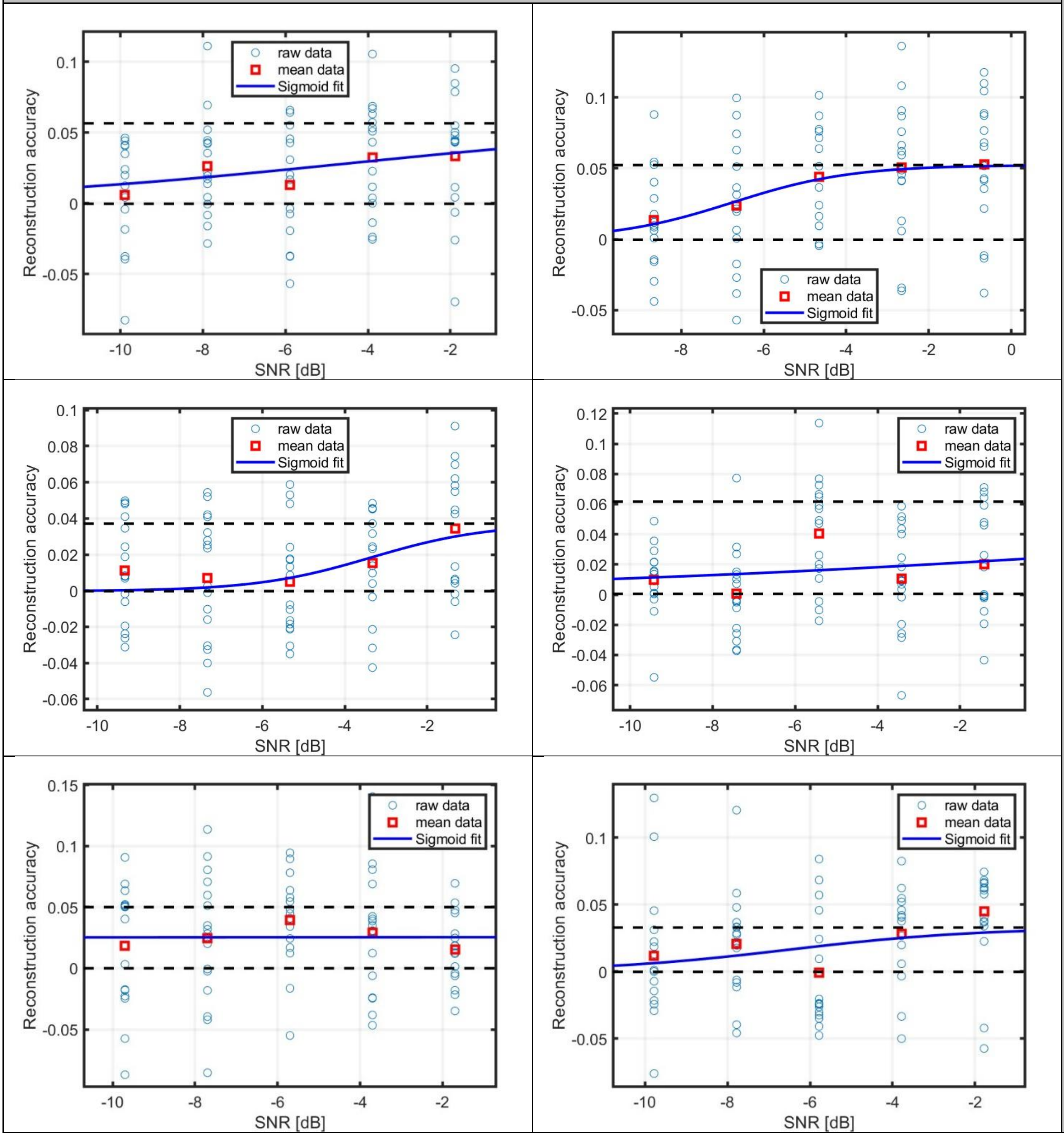
raw data
mean data
Sigmoid fit
Reconstruction accuracy
SNR [dB]

EarT

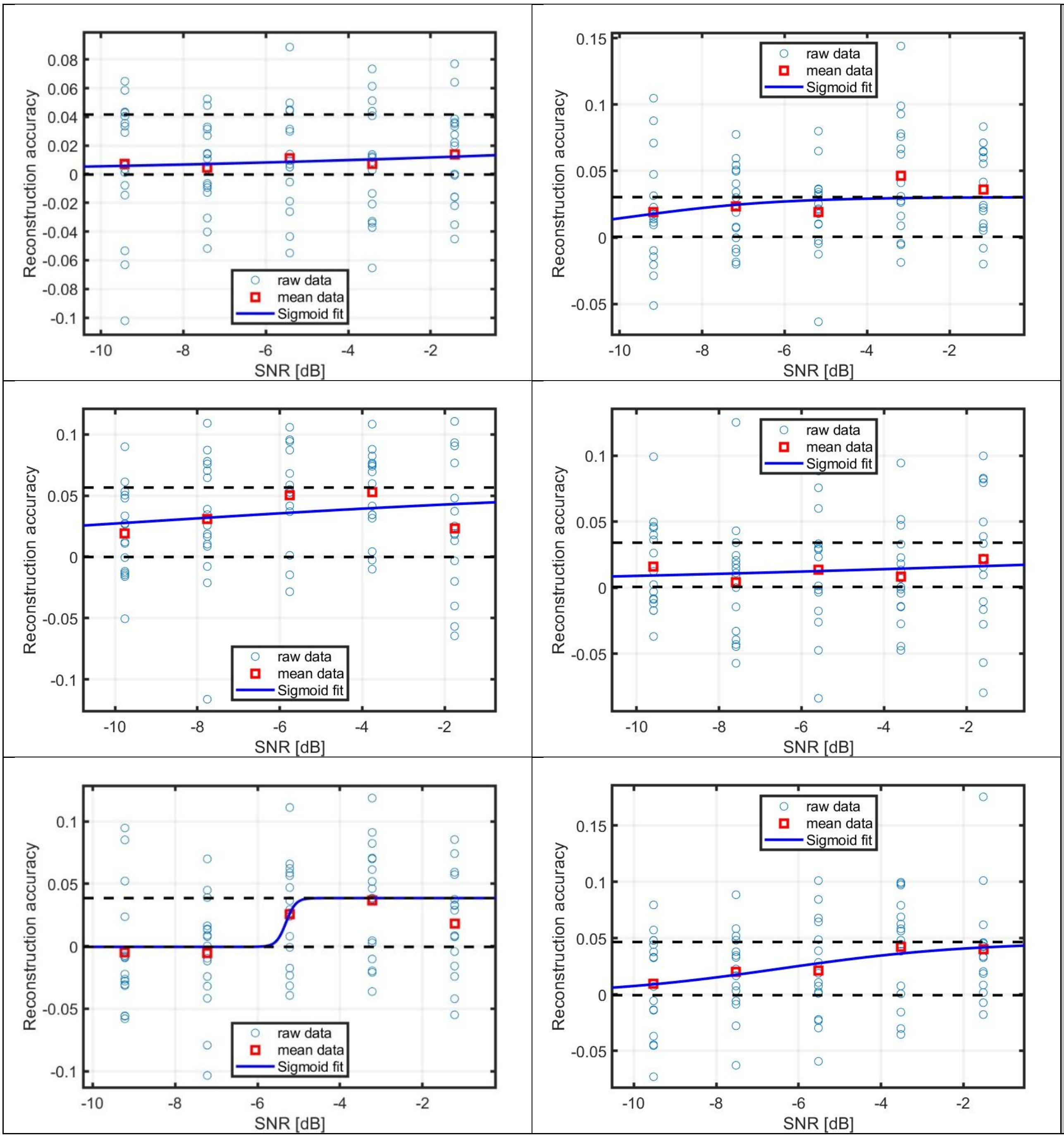
Reconstruction accuracy
SNR [dB]
raw data
mean data
Sigmoid fit

EarT

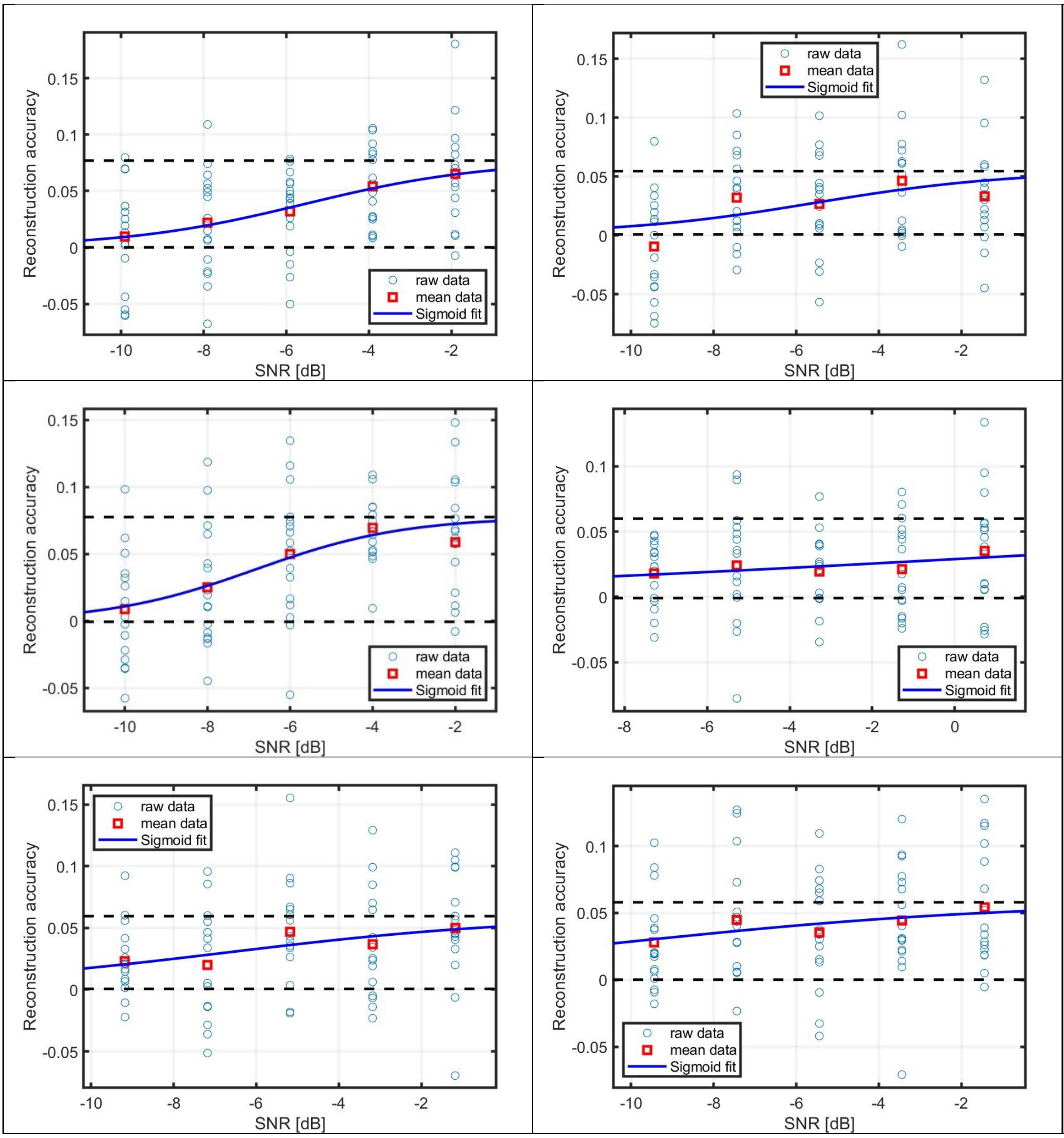
Reconstruction accuracy
SNR [dB]
raw data
mean data
Sigmoid fit

EarT

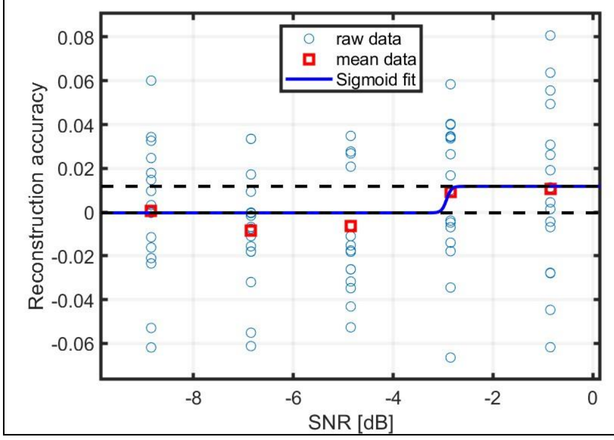


EarT

AEarT

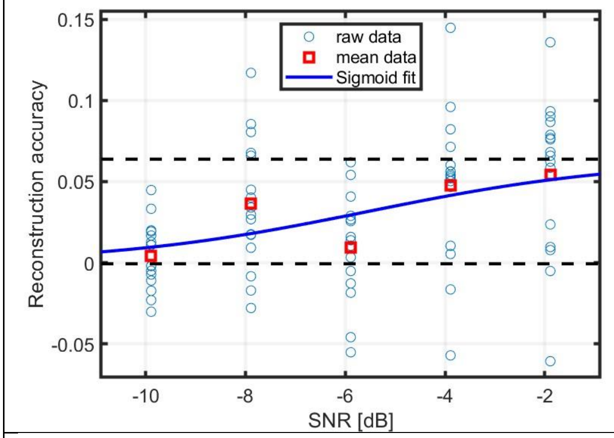


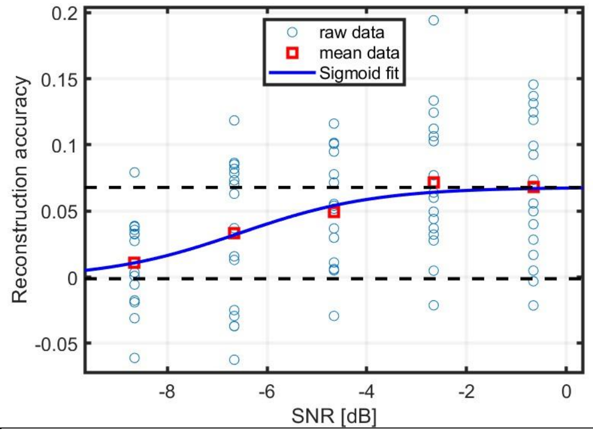


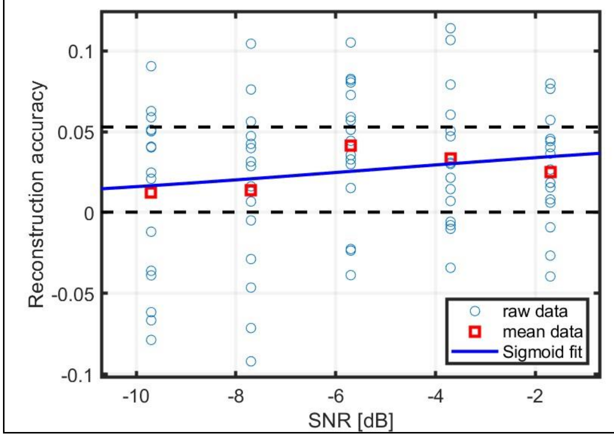


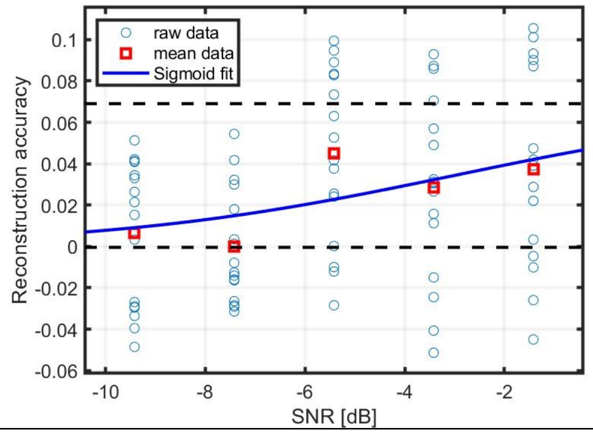


AEarT

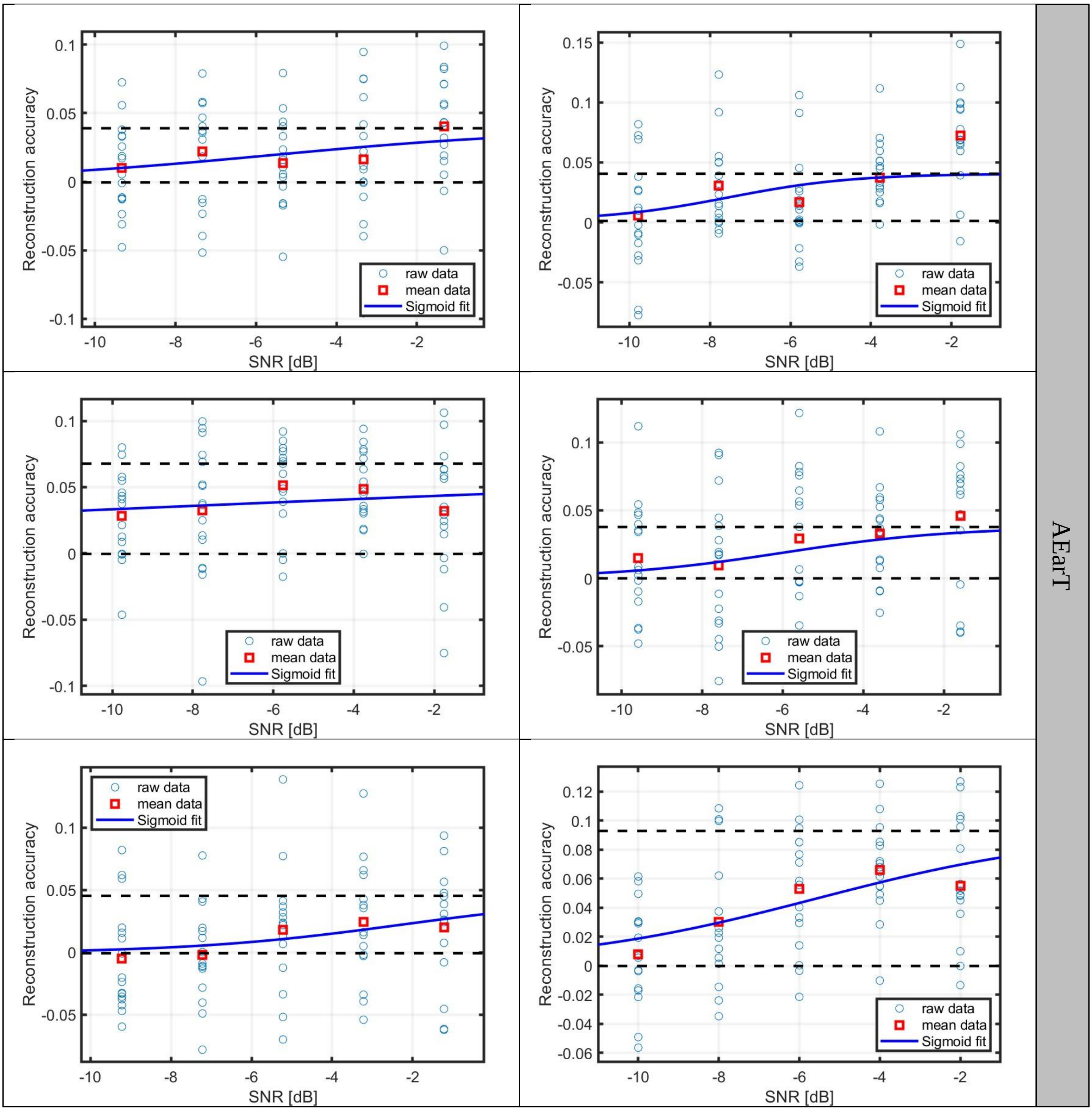

Reconstruction accuracy
SNR [dB]
raw data
mean data
Sigmoid fit
AEarT

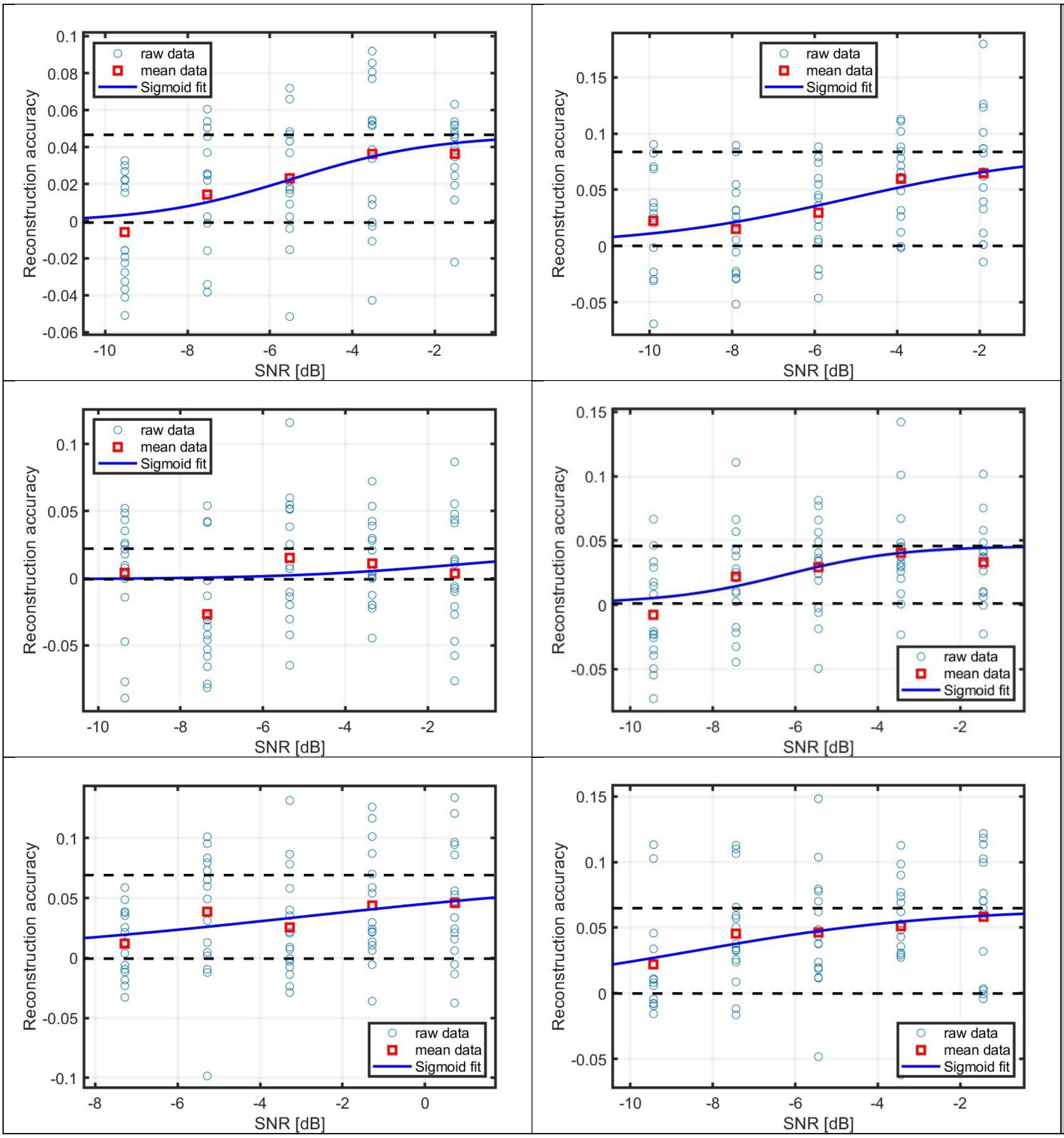
raw data
mean data
Sigmoid fit
Reconstruction accuracy
SNR [dB]

AEarT

AEarT

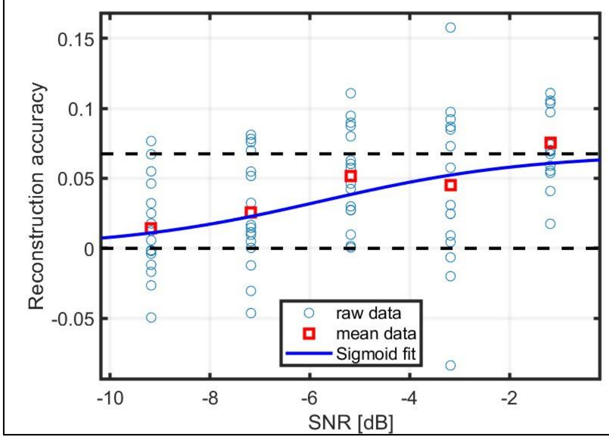
Reconstruction accuracy
SNR [dB]
raw data
mean data
Sigmoid fit

MT

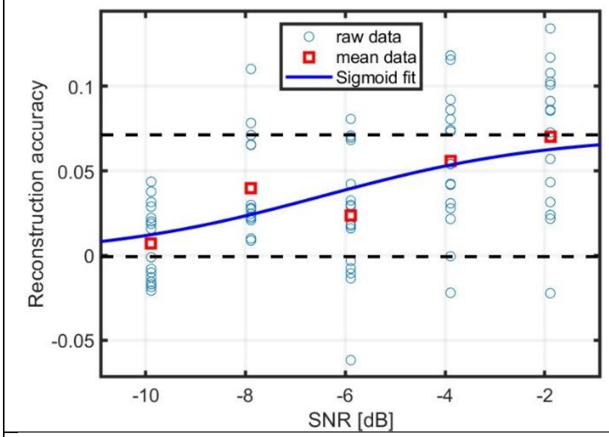
Reconstruction accuracy
SNR [dB]
raw data
mean data
Sigmoid fit

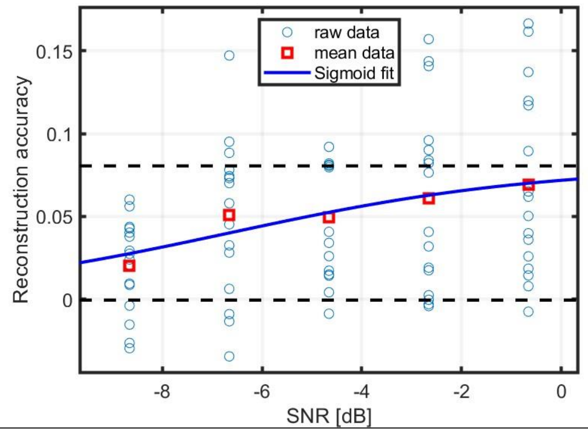
Reconstruction accuracy
SNR [dB]
raw data
mean data
Sigmoid fit

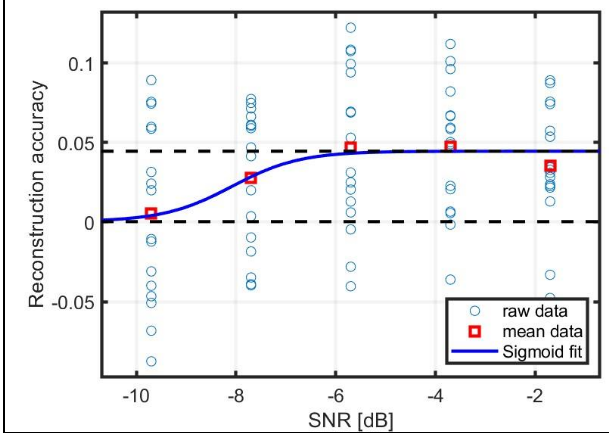
Reconstruction accuracy
SNR [dB]
raw data
mean data
Sigmoid fit

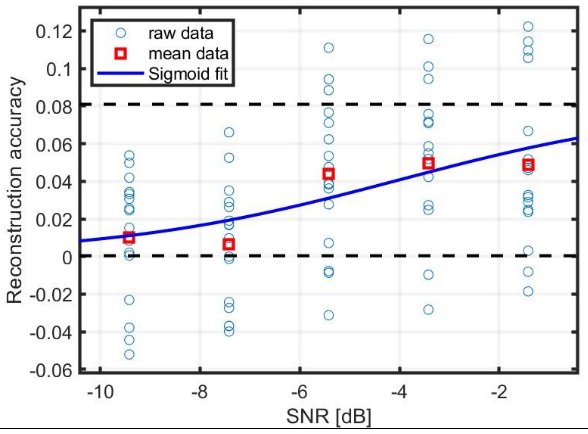
Reconstruction accuracy
SNR [dB]
raw data
mean data
Sigmoid fit

M

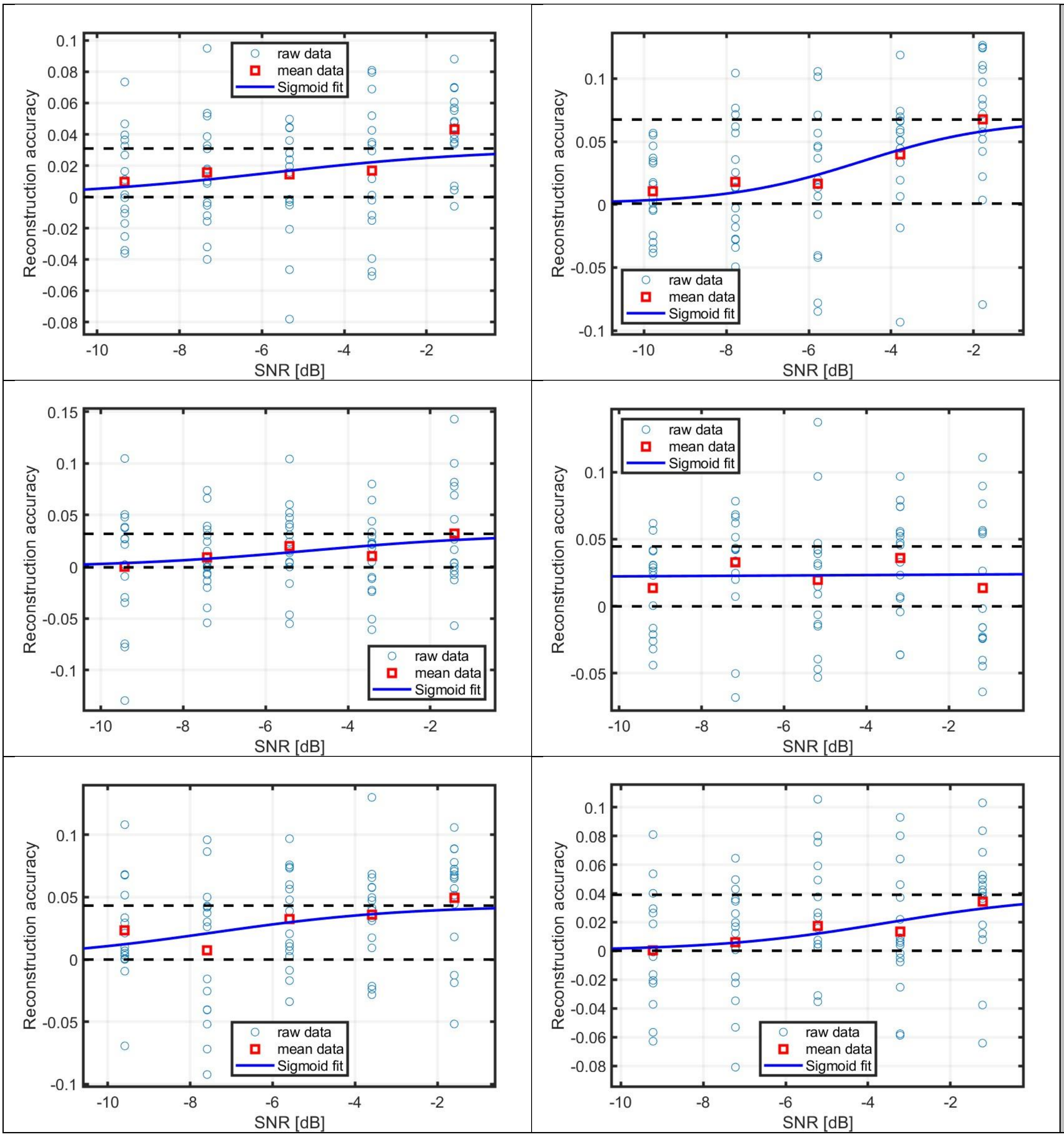
raw data
mean data
Sigmoid fit
Reconstruction accuracy
SNR [dB]

M

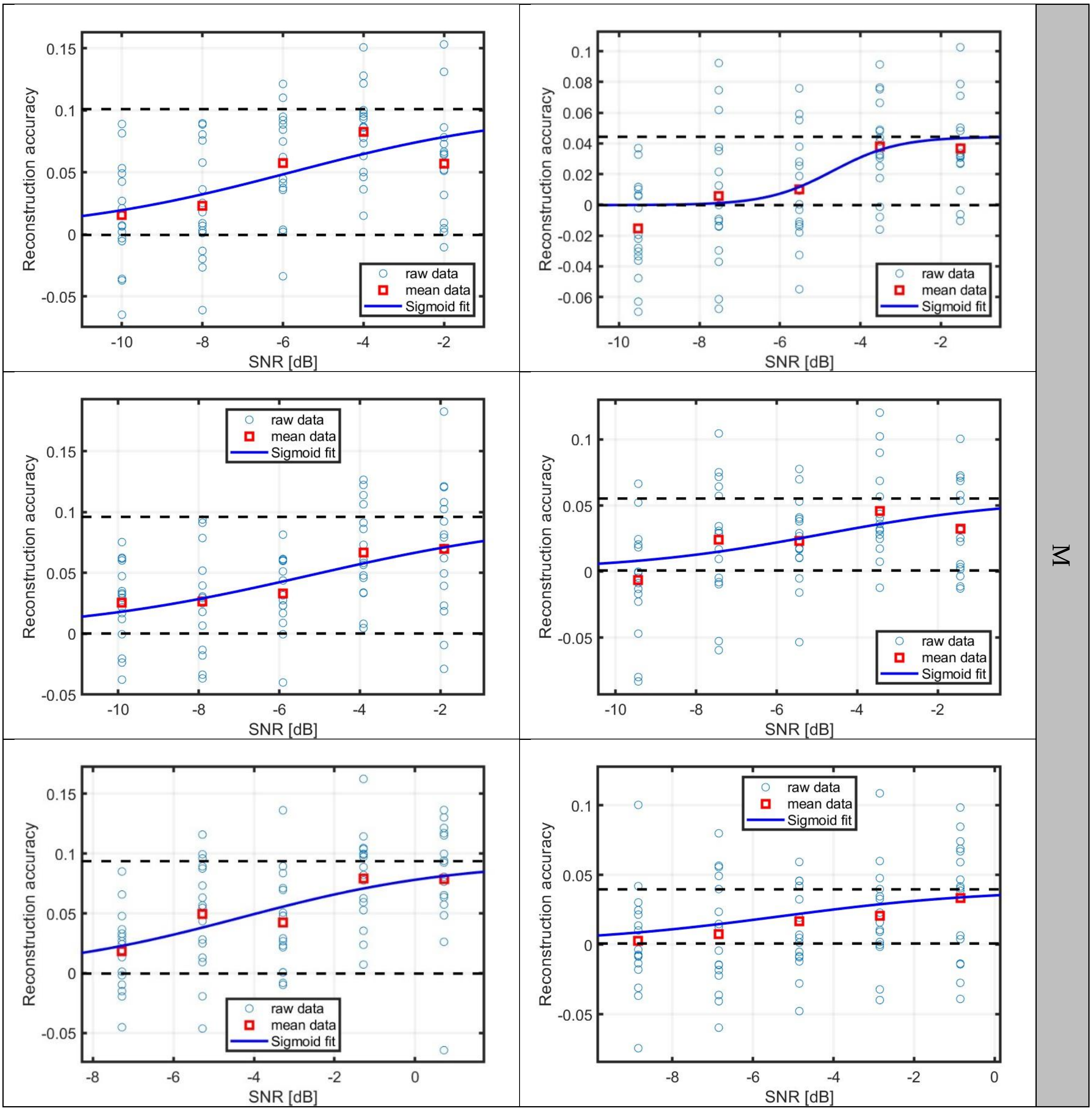
Reconstruction accuracy
SNR [dB]
raw data
mean data
Sigmoid fit
M

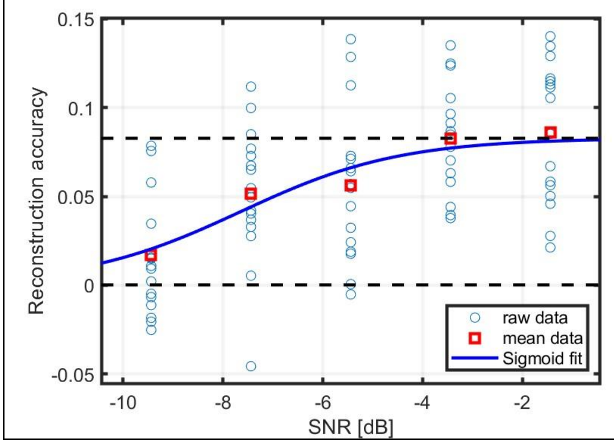


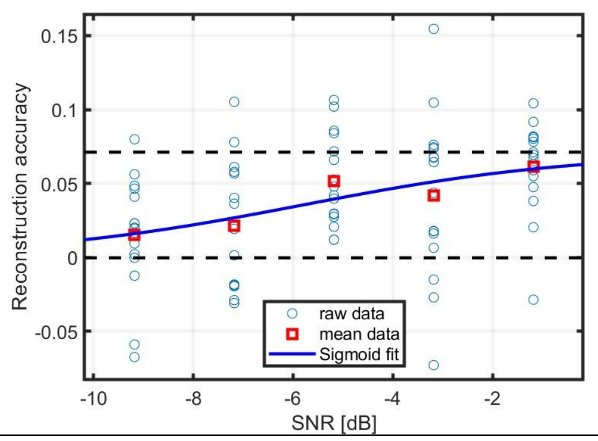


M

EarMT

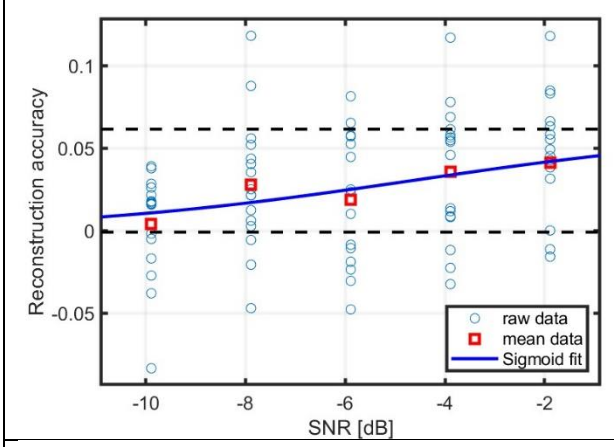


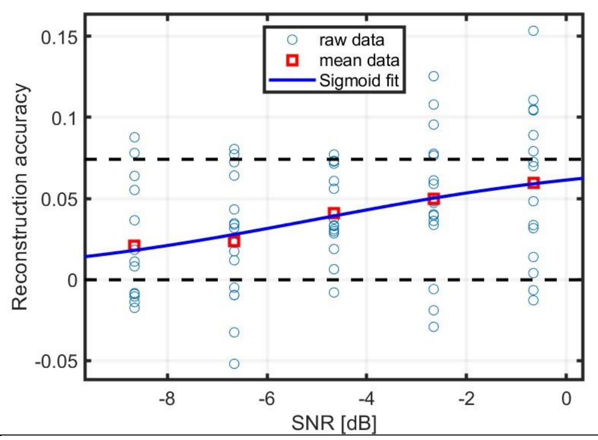


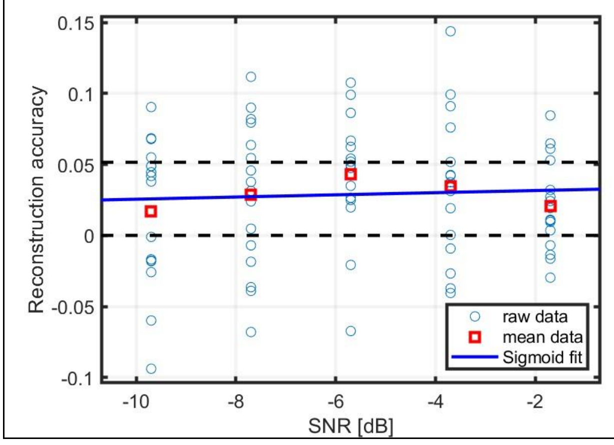


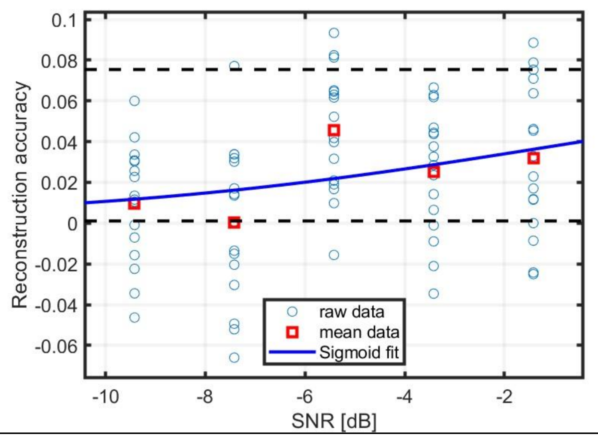


EarMT

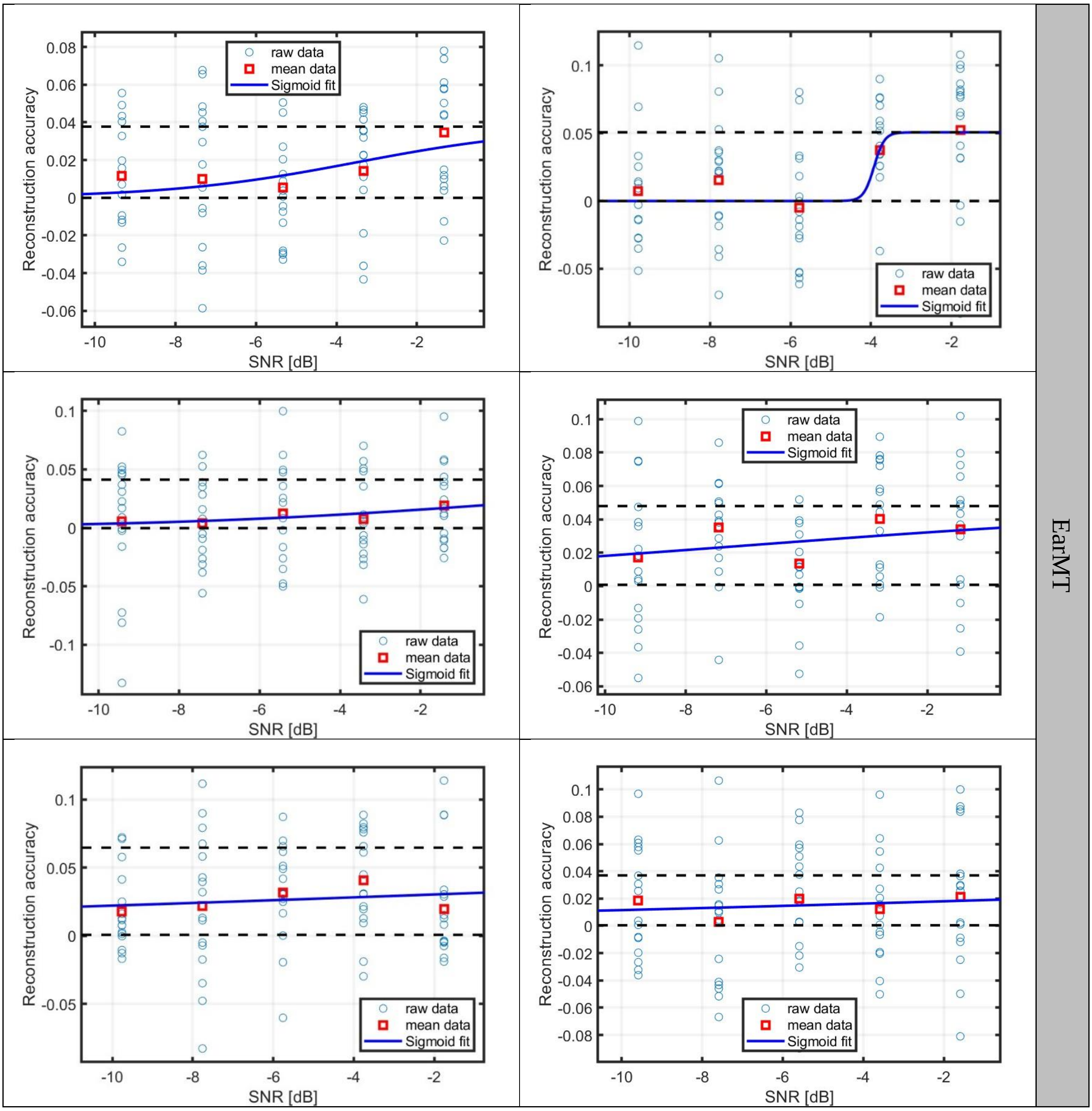
EarMT
raw data
mean data
Sigmoid fit
Reconstruction accuracy
SNR [dB]

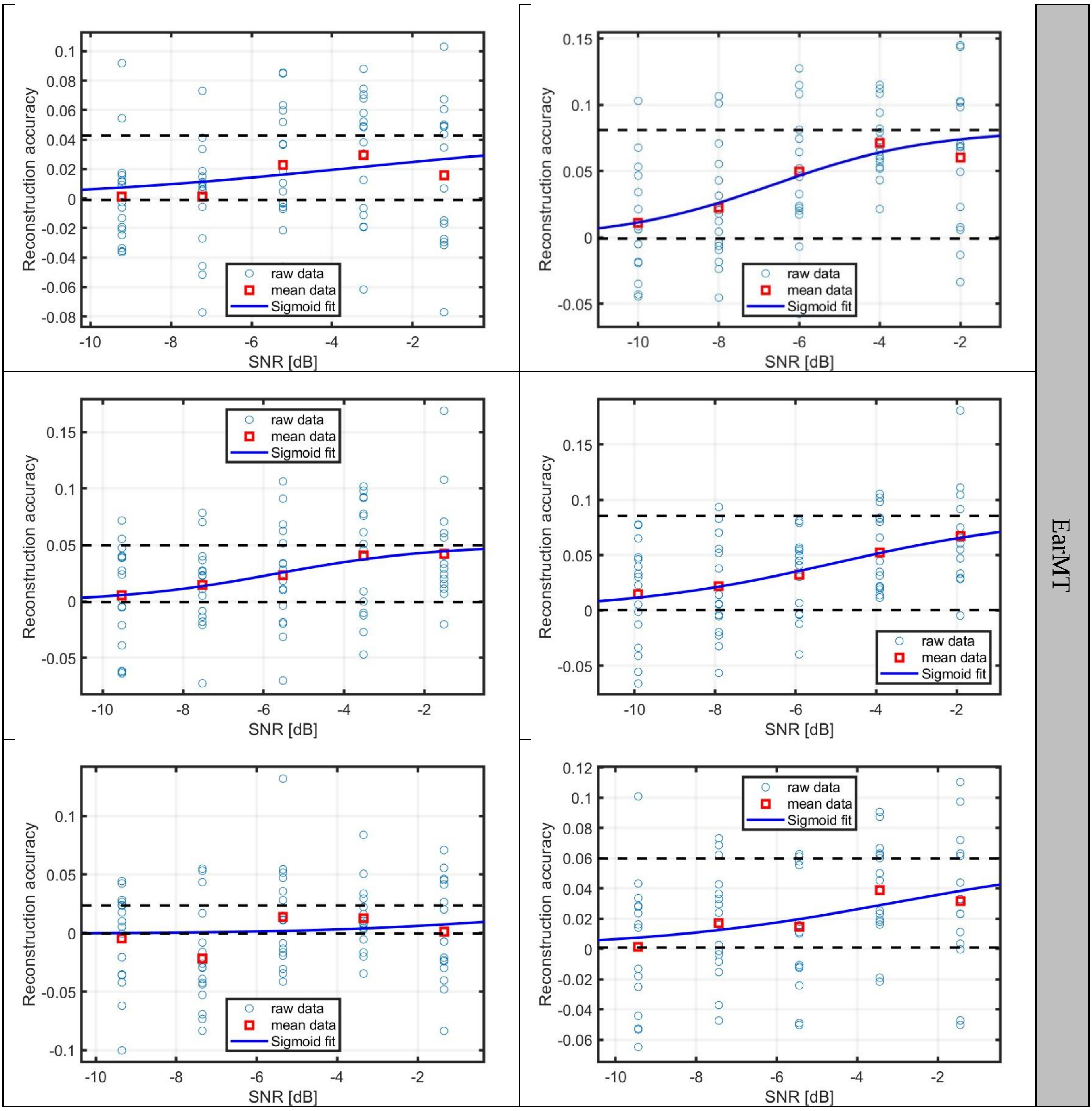

Reconstruction accuracy
SNR [dB]
raw data
mean data
Sigmoid fit
EarMT

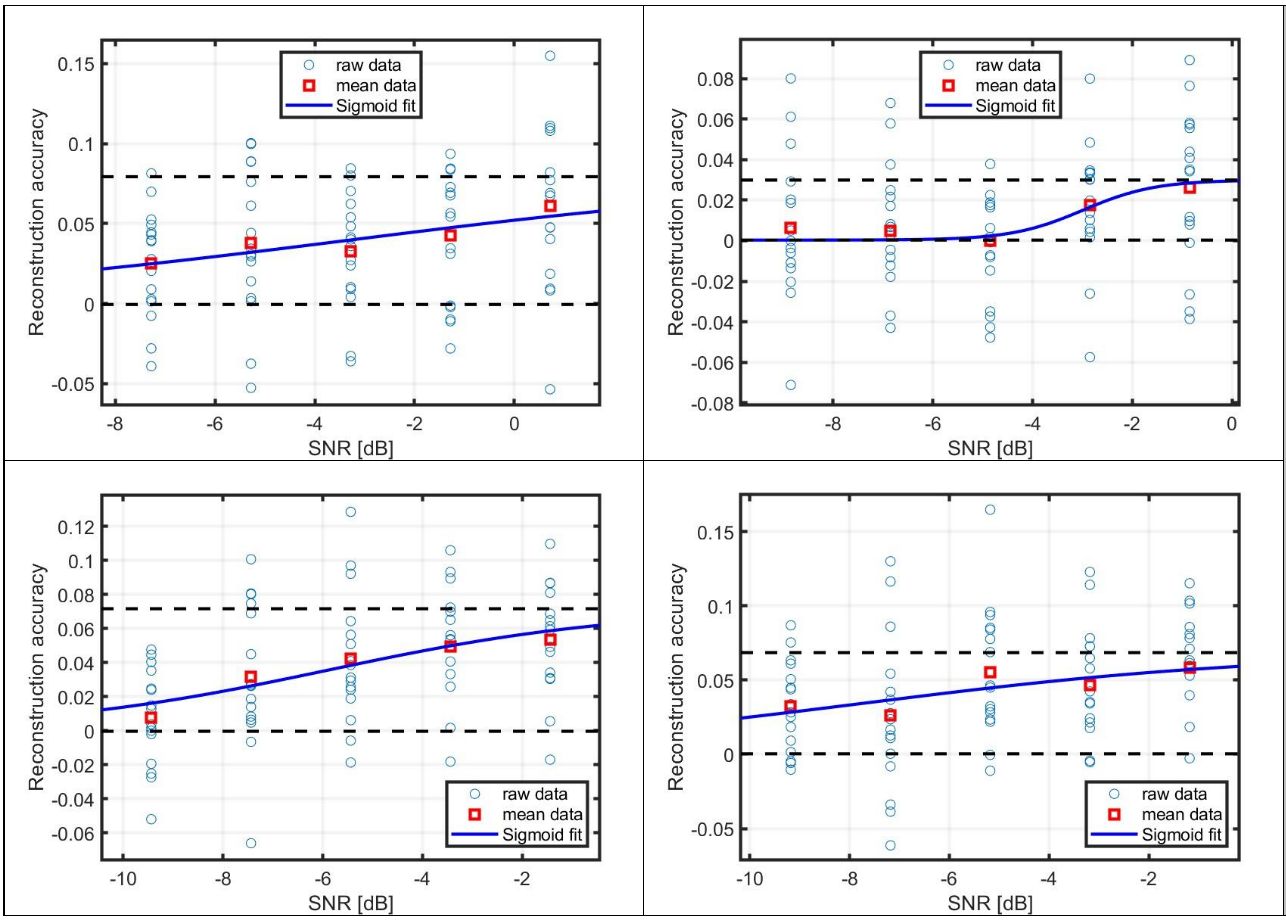


EarMT

LAEarMT

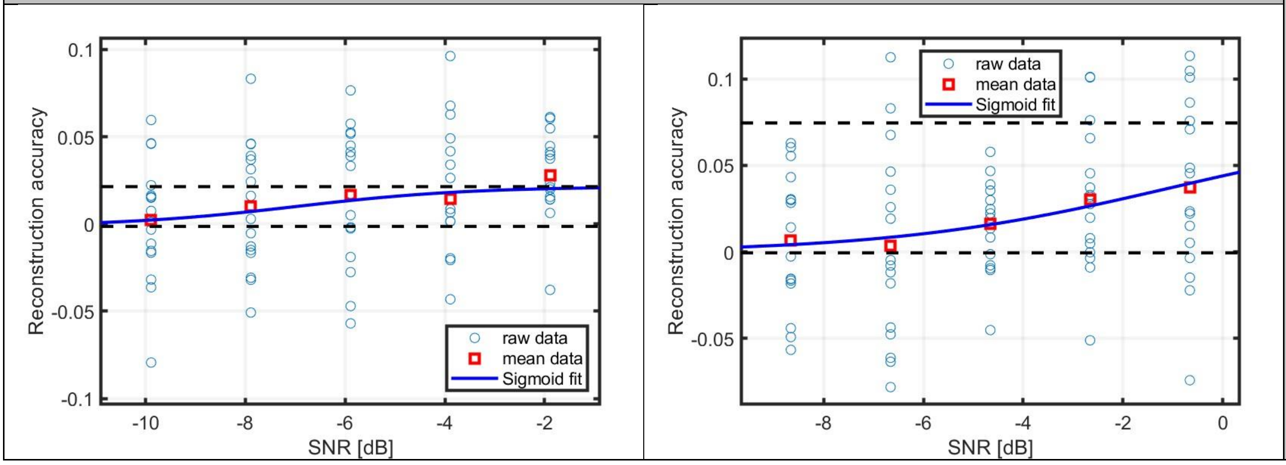


LaEarMT

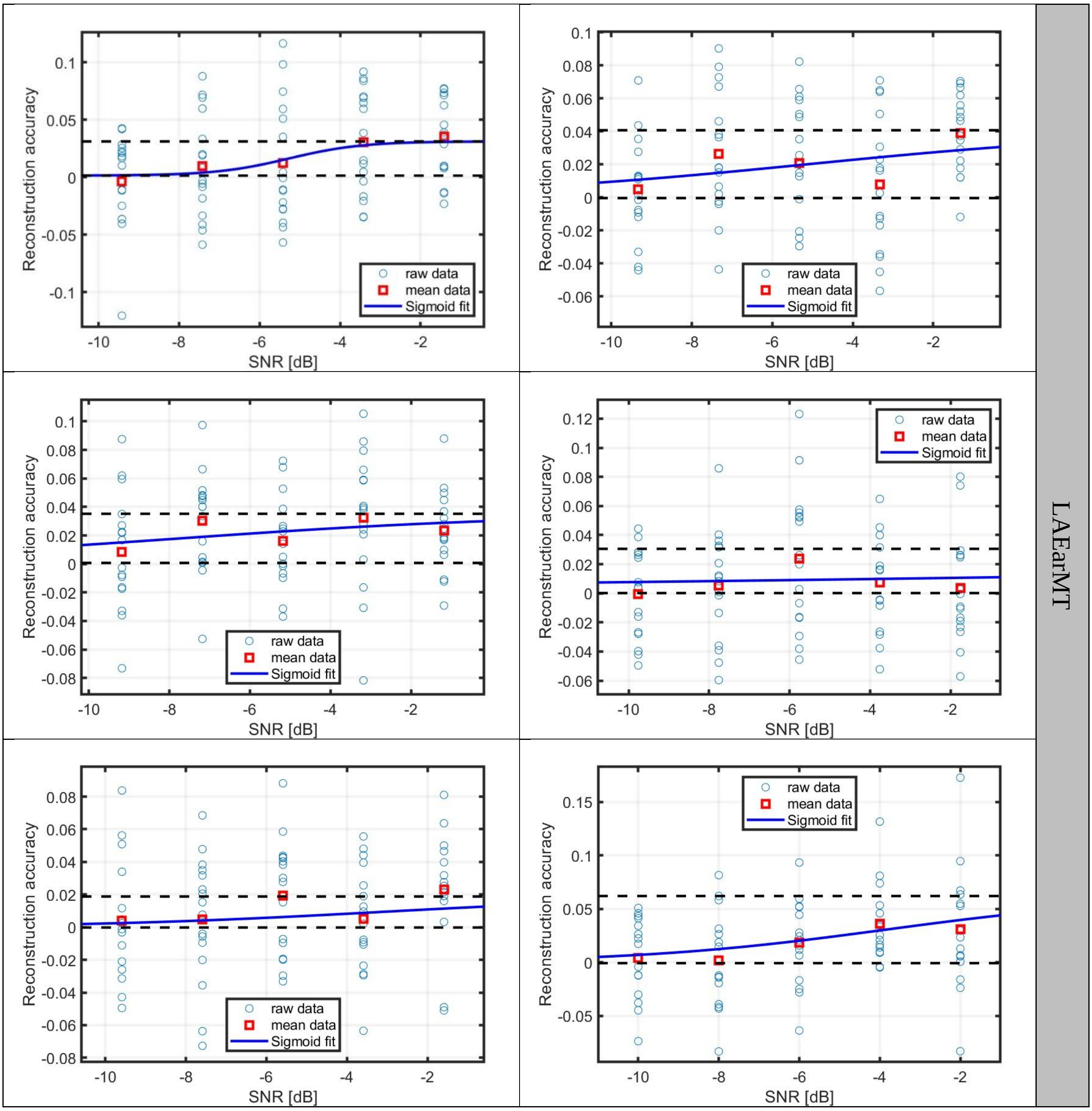
Reconstruction accuracy
SNR [dB]
raw data
mean data
Sigmoid fit
LAEarMT

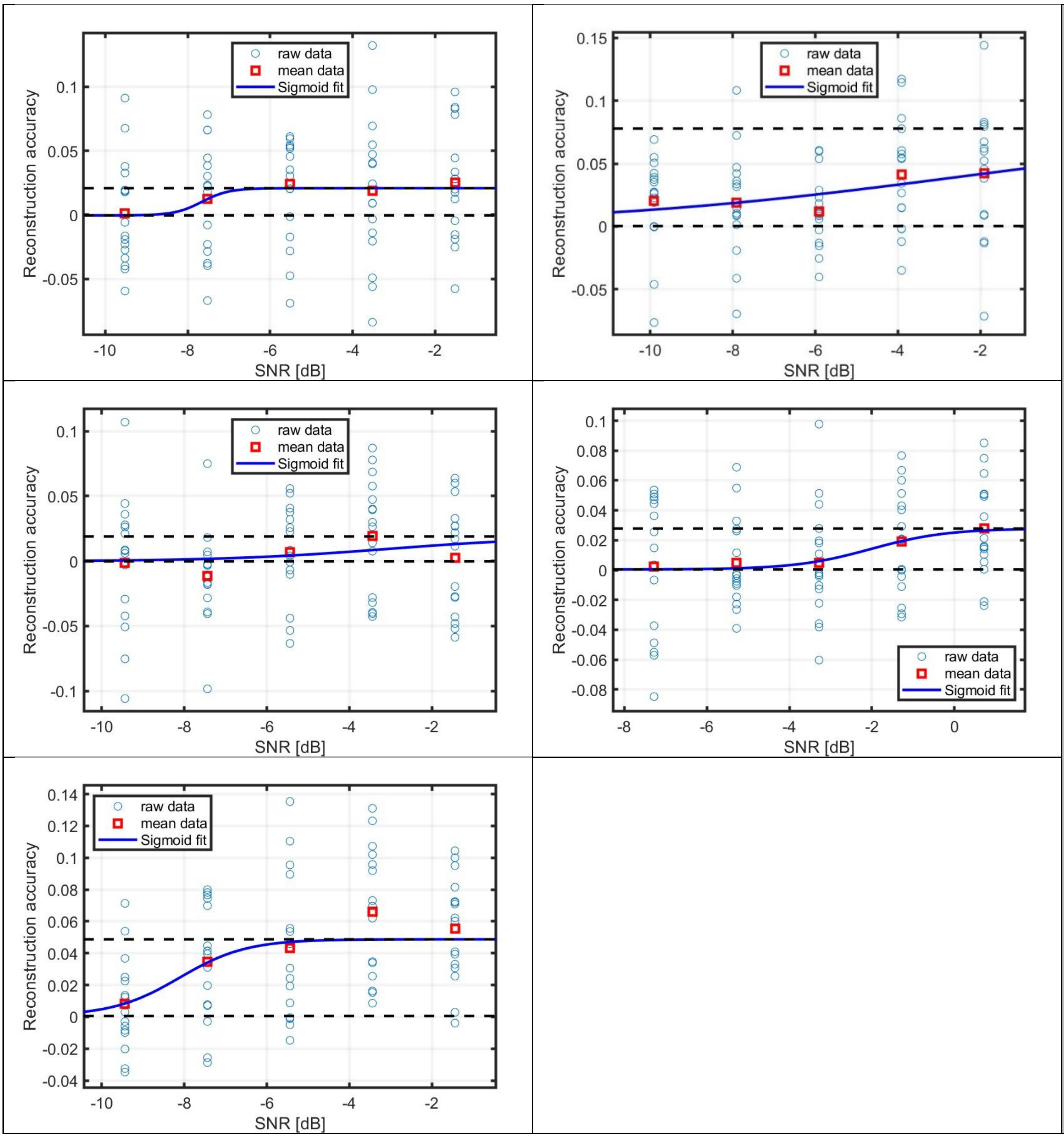
raw data
mean data
Sigmoid fit
Reconstruction accuracy
SNR [dB]

LaEarMT

RAEarMT

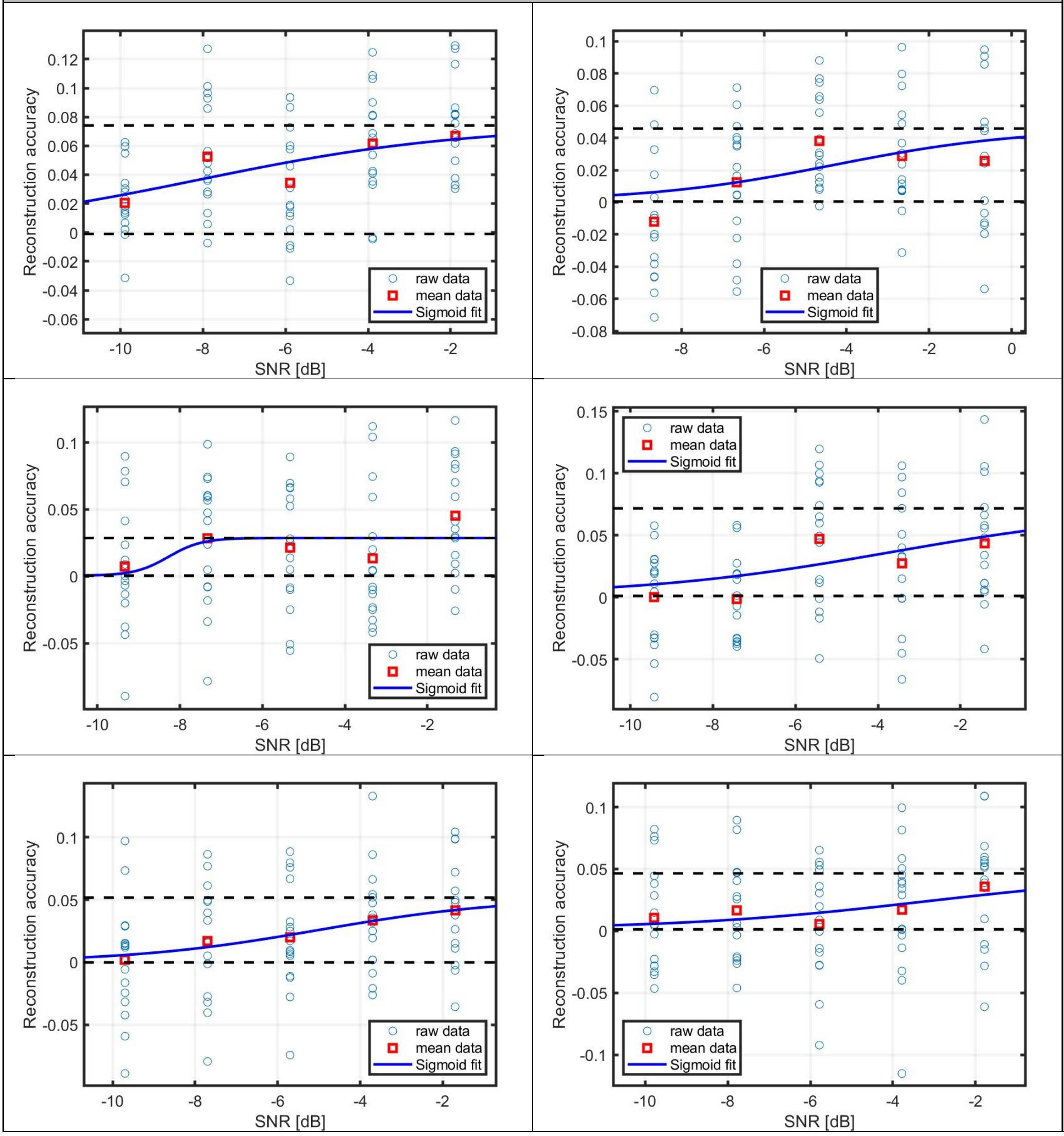


RAEarMT

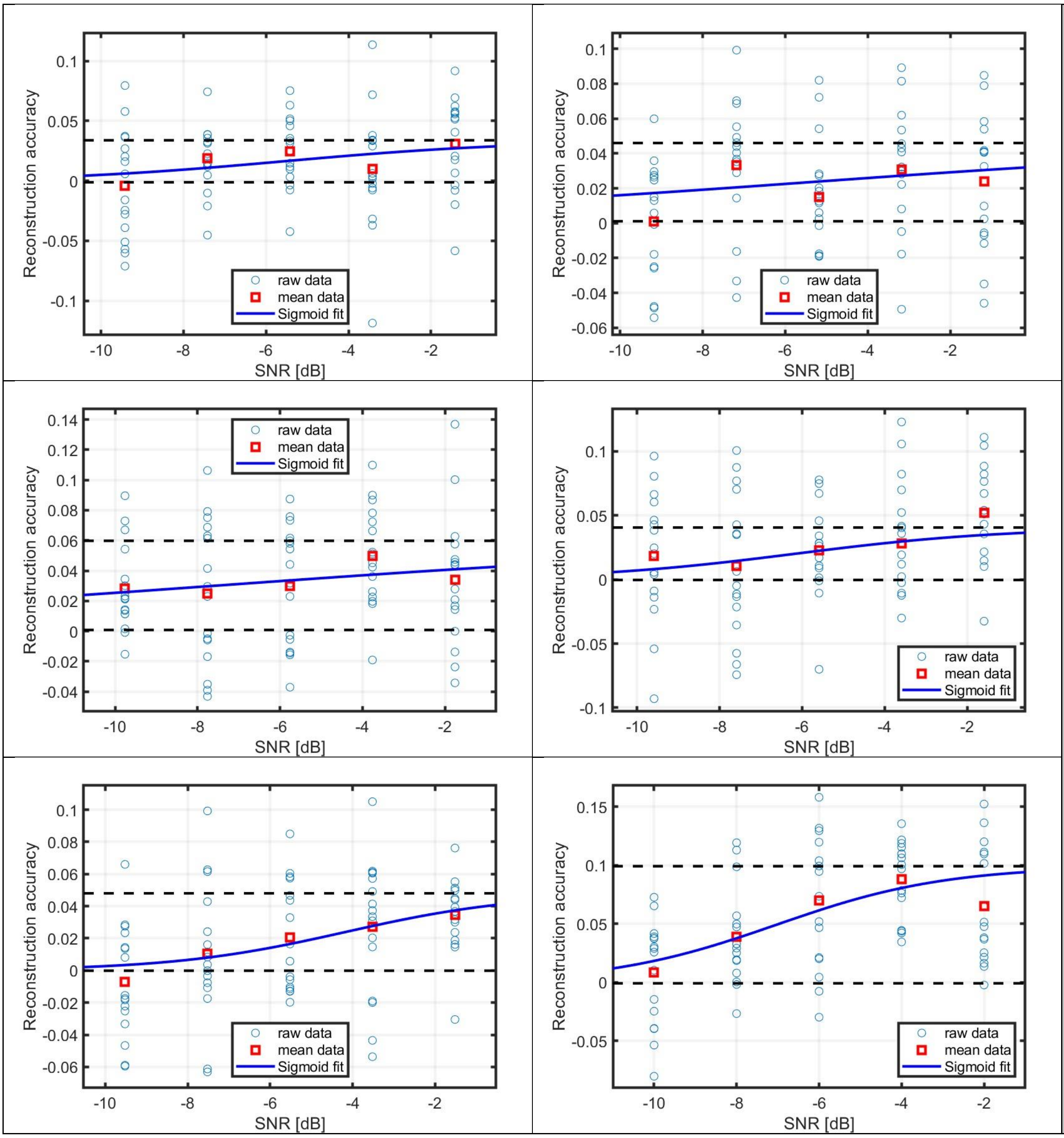
Reconstruction accuracy
SNR [dB]
raw data
mean data
Sigmoid fit

RAEarMT

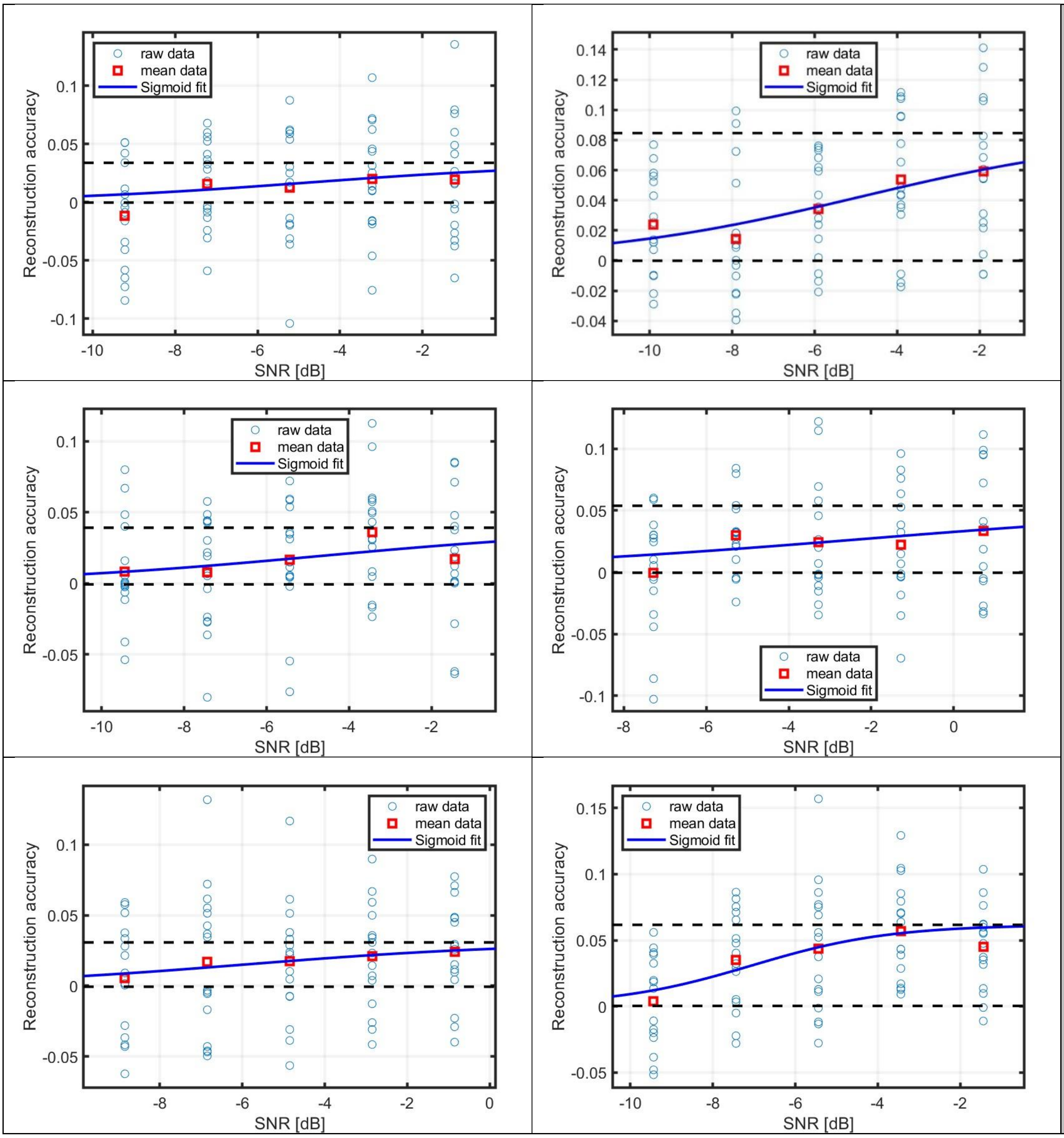
raw data
mean data
Sigmoid fit
Reconstruction accuracy
SNR [dB]

RAEarMT

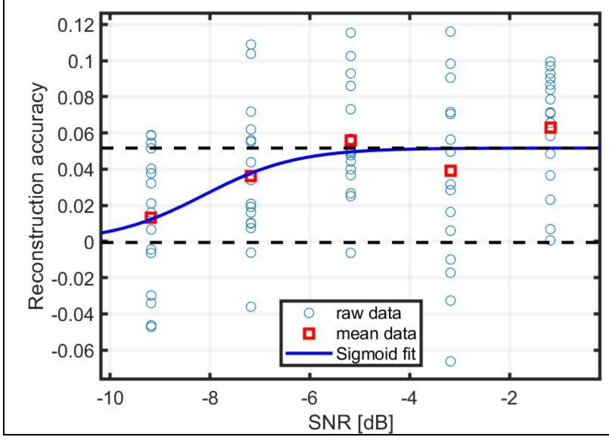


RAEarMT

AEarMT

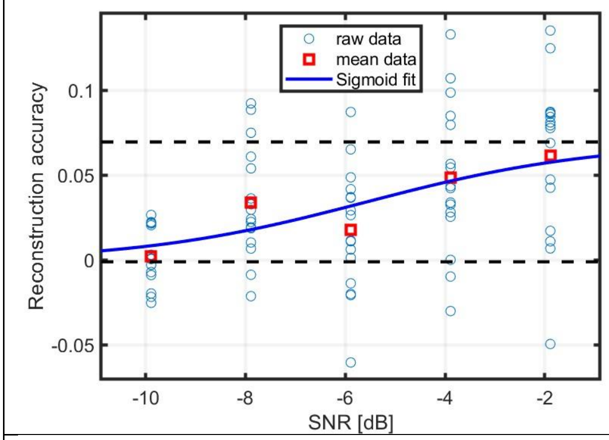


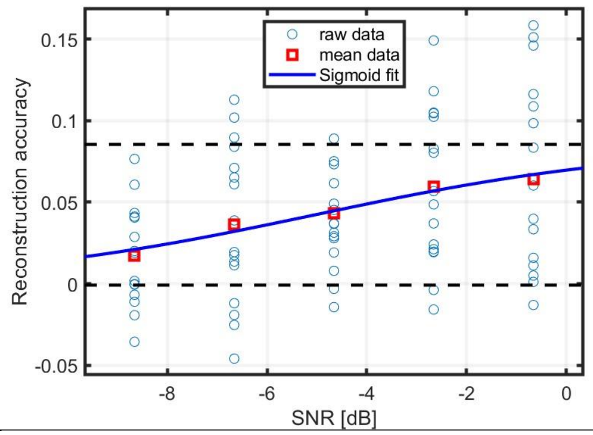


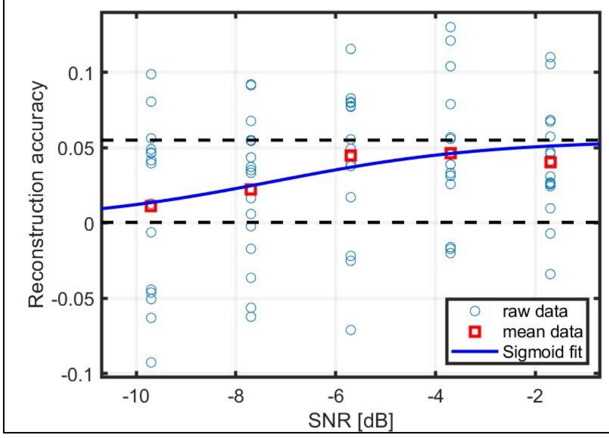


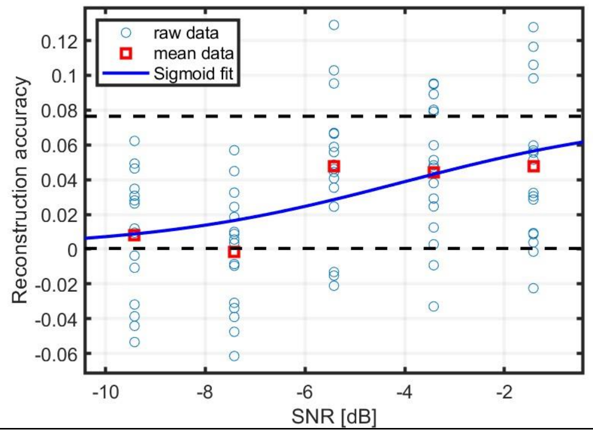


AEarMT

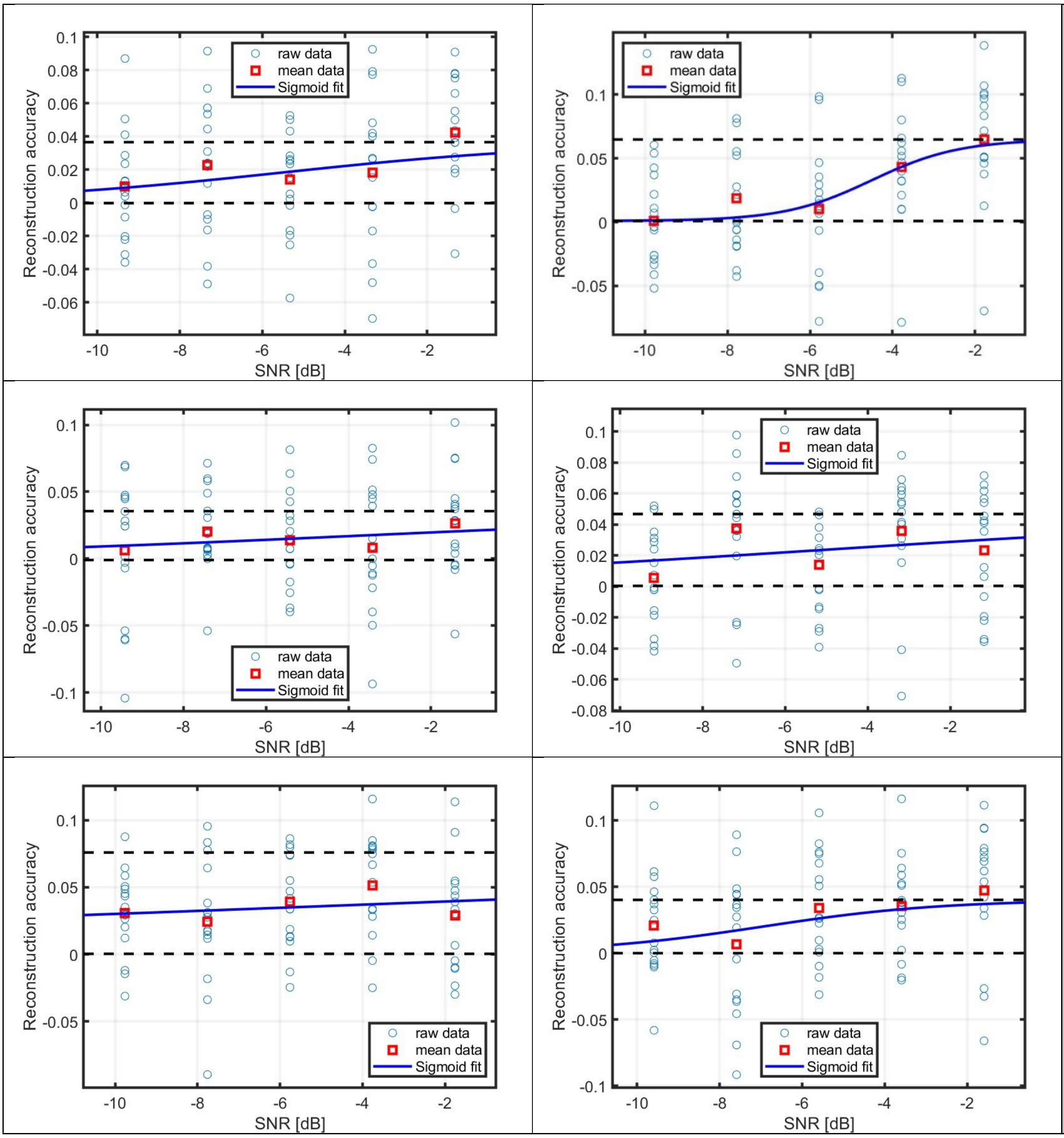
raw data
mean data
Sigmoid fit
Reconstruction accuracy
SNR [dB]

AEarMT

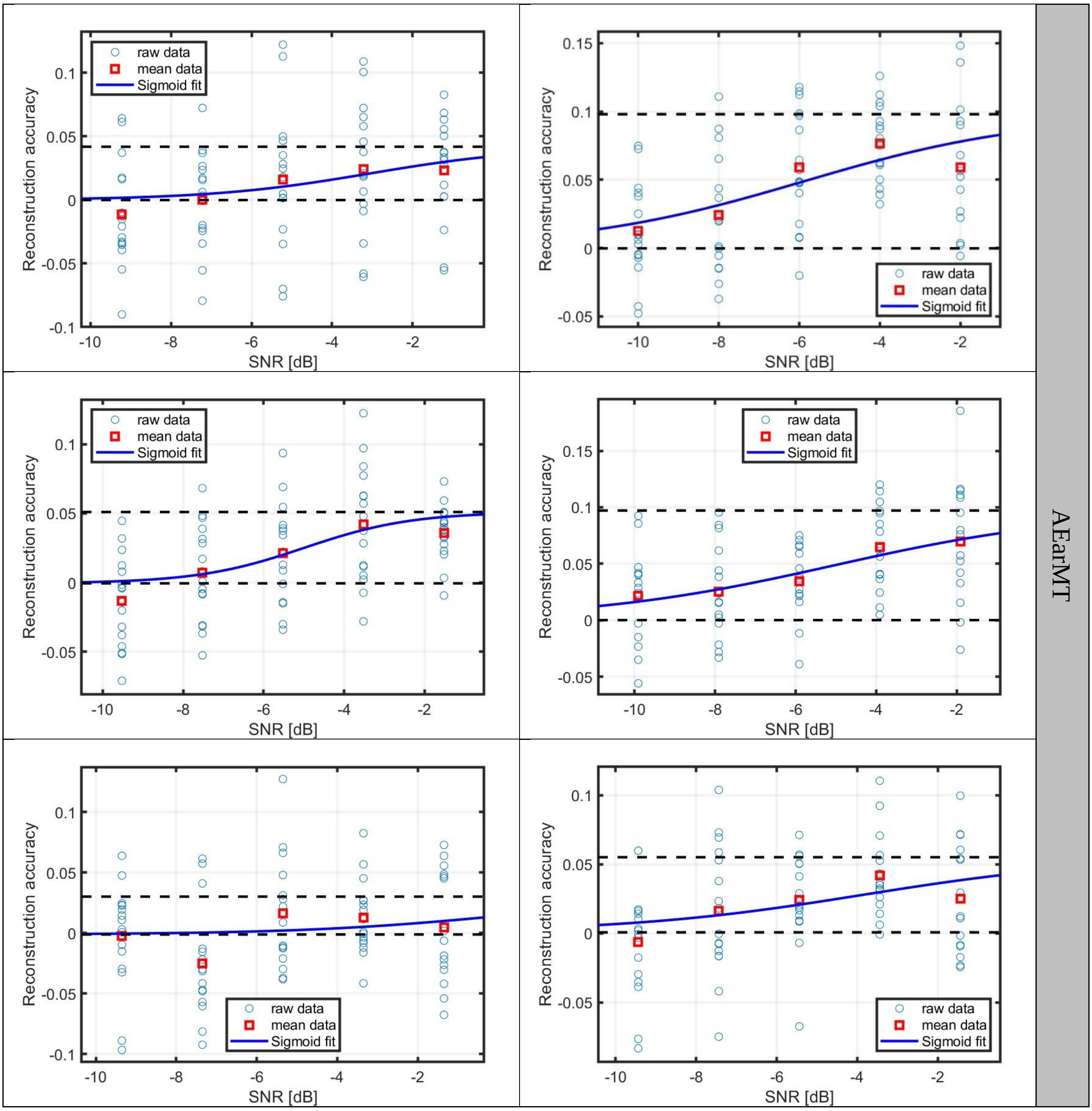
raw data
mean data
Sigmoid fit
Reconstruction accuracy
SNR [dB]
AEarMT

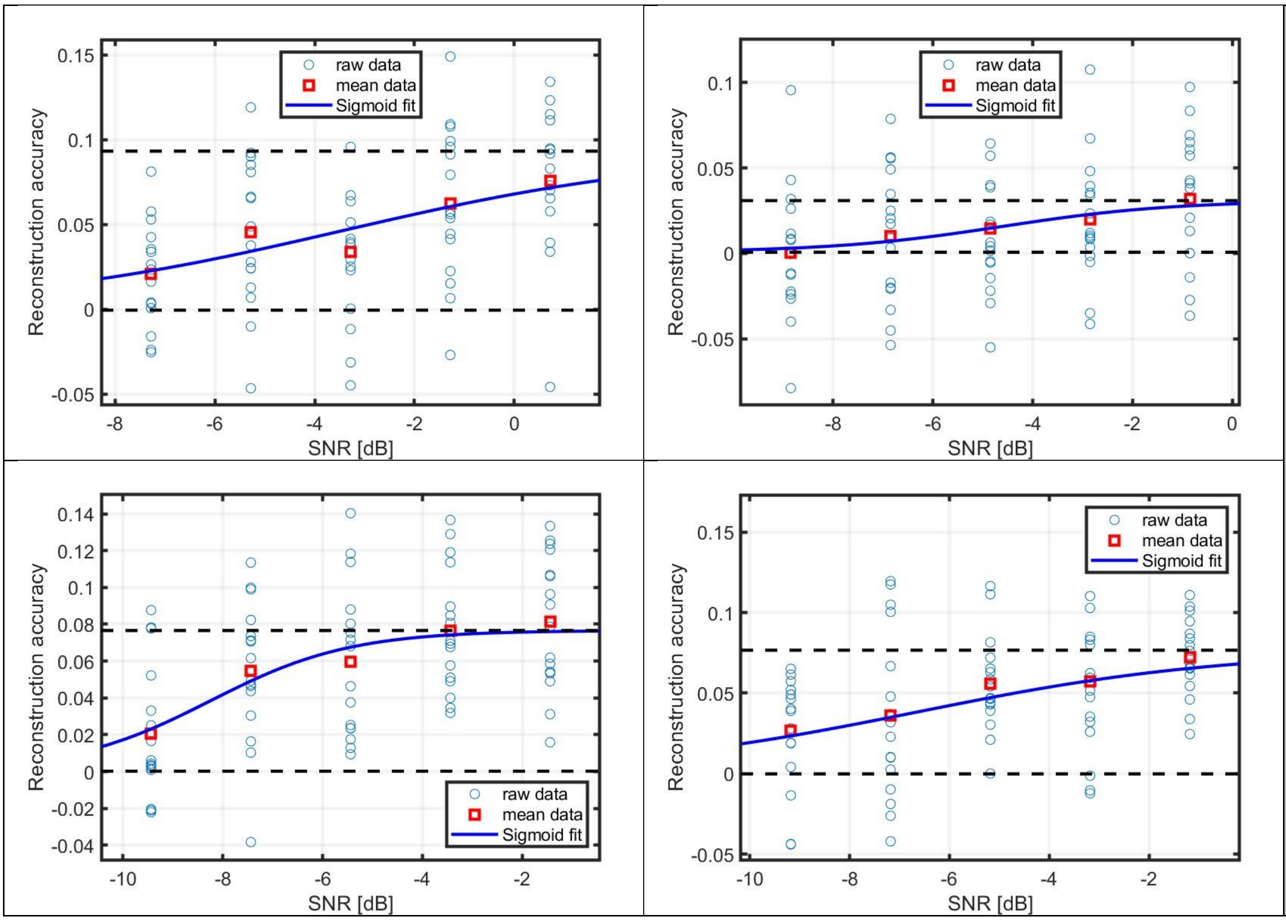


AEarMT

Scalp

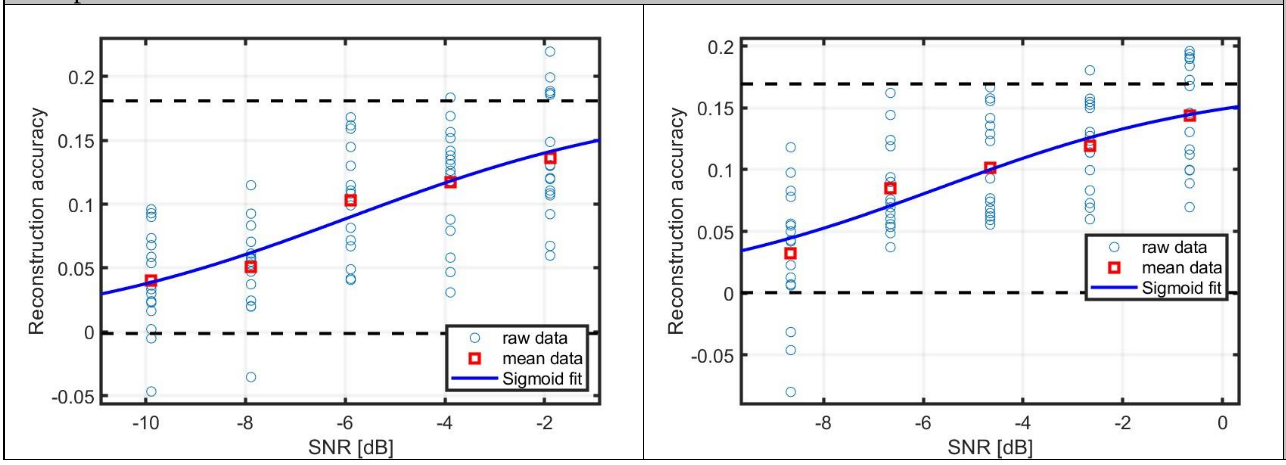


Scalp

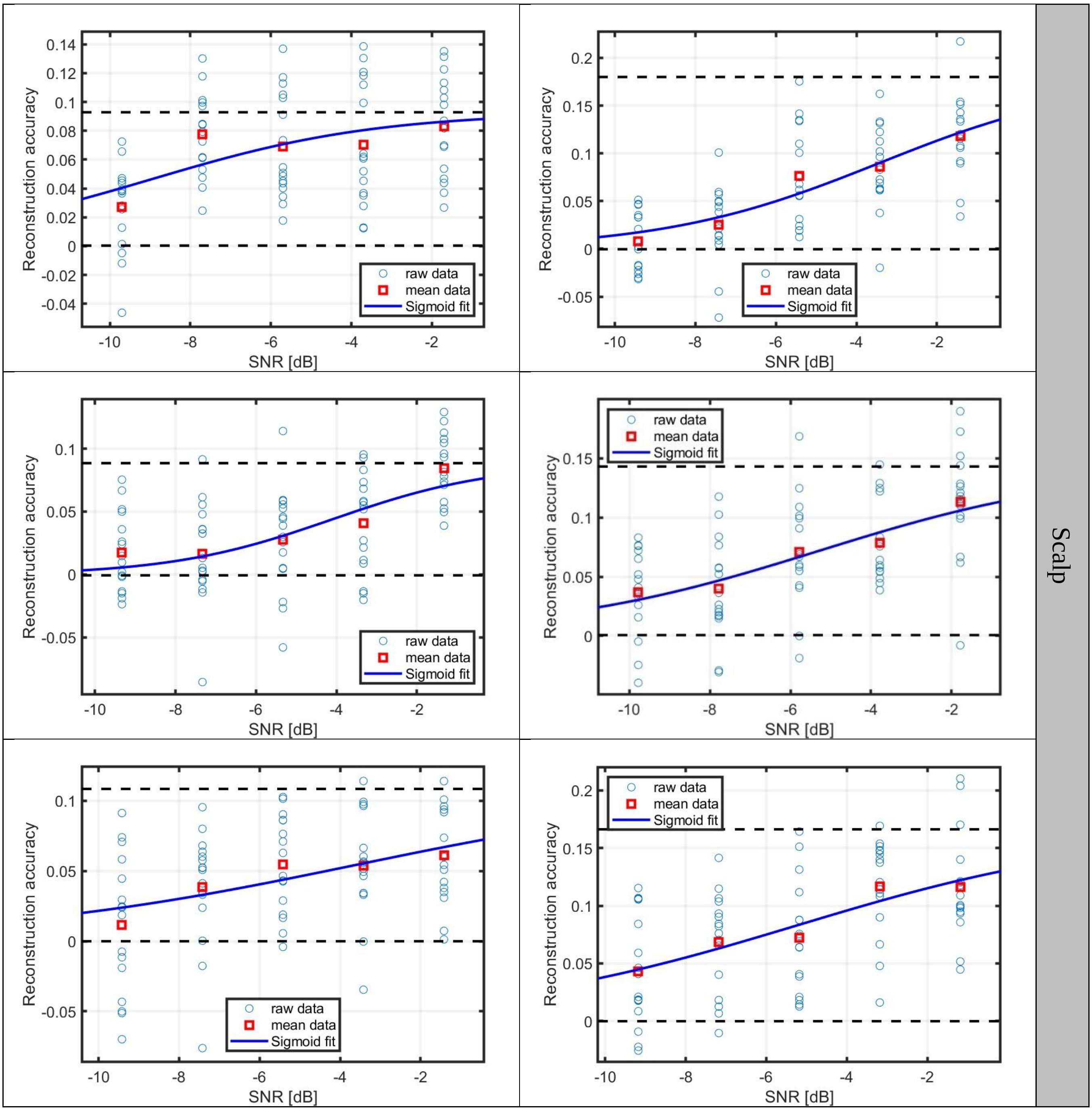
Reconstruction accuracy
SNR [dB]
raw data
mean data
Sigmoid fit
Scalp

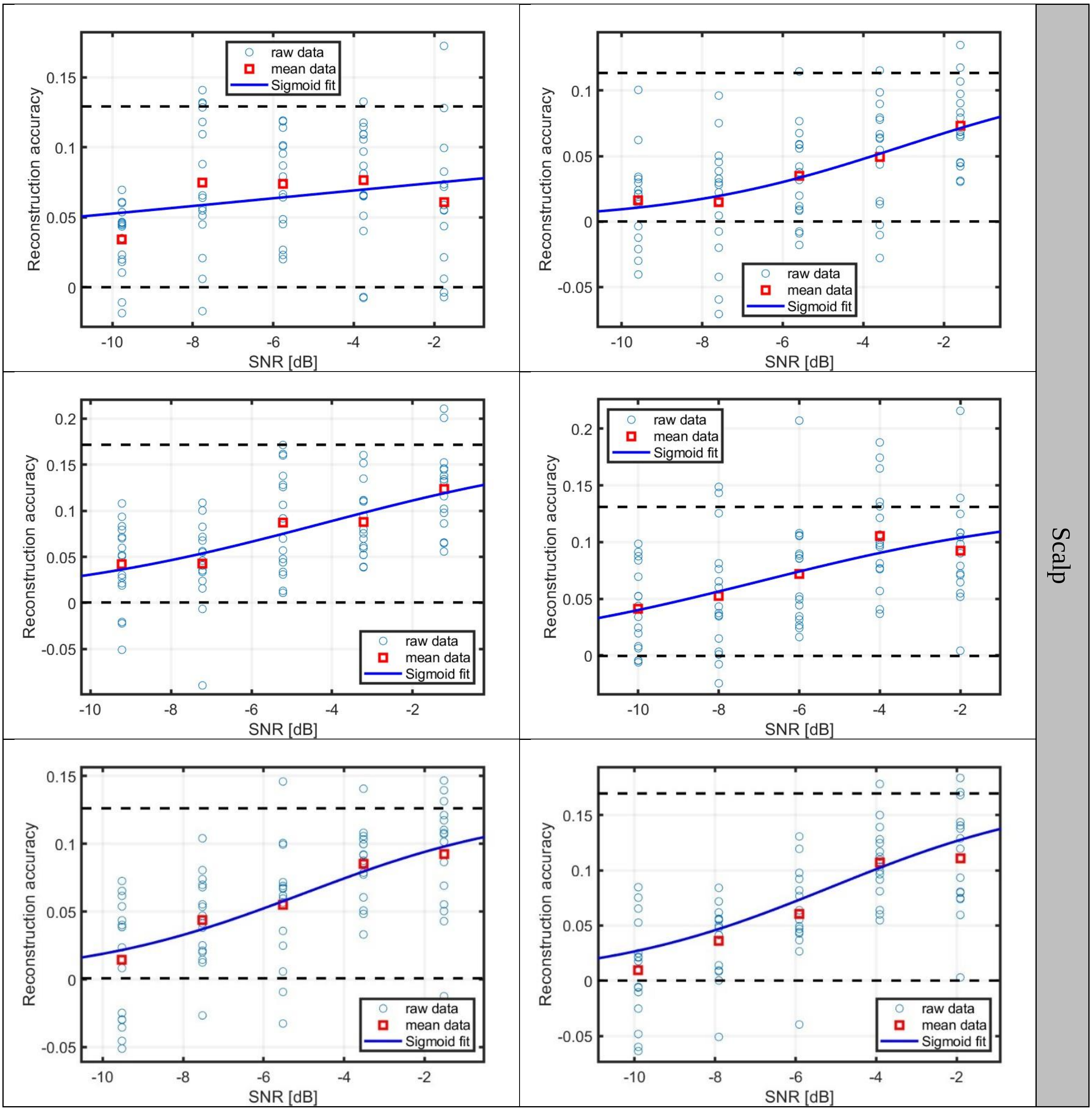
raw data
mean data
Sigmoid fit
Reconstruction accuracy
SNR [dB]
Scalp

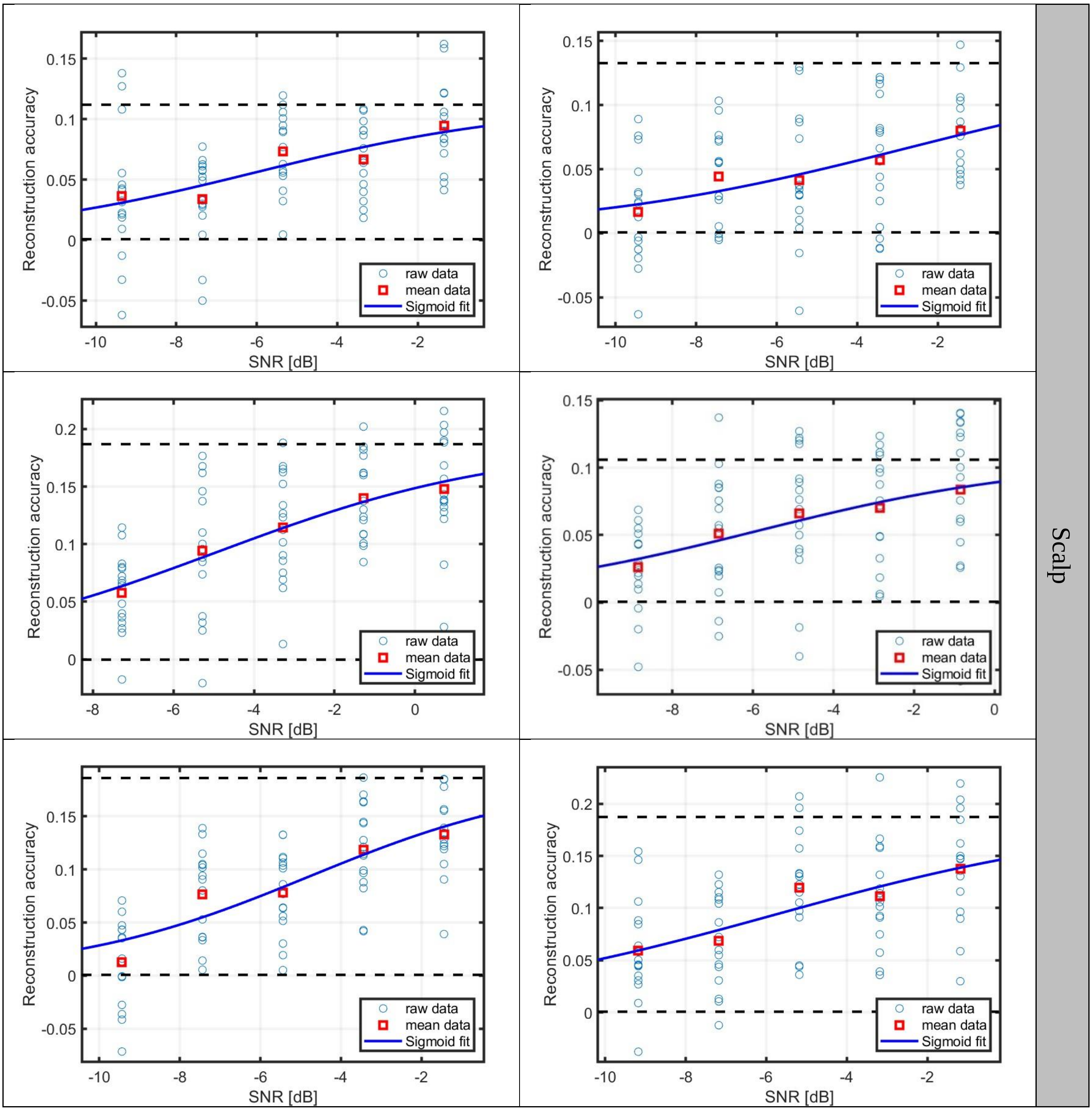
Reconstruction accuracy
SNR [dB]
raw data
mean data
Sigmoid fit
Scalp
